# Locus coeruleus integrity predicts ease of attaining and maintaining neural states of high attentiveness

**DOI:** 10.1101/2022.03.07.483289

**Authors:** Sana Hussain, Isaac Menchaca, Mahsa Alizadeh Shalchy, Kimia Yaghoubi, Jason Langley, Aaron R. Seitz, Xiaoping P. Hu, Megan A. K. Peters

## Abstract

The locus coeruleus (LC), a small subcortical structure in the brainstem, is the brain’s principal source of norepinephrine. It plays a primary role in regulating stress, the sleep-wake cycle, and attention, and its degradation is associated with aging and neurodegenerative diseases associated with cognitive deficits (e.g., Parkinson’s, Alzheimer’s). Yet precisely how norepinephrine drives brain networks to support healthy cognitive function remains poorly understood – partly because LC’s small size makes it difficult to study noninvasively in humans. Here, we characterized LC’s influence on brain dynamics using a hidden Markov model fitted to functional neuroimaging data from healthy young adults across four attention-related brain networks and LC. We modulated LC activity using a behavioral paradigm and measured individual differences in LC magnetization transfer contrast. The model revealed five hidden states, including a stable state dominated by salience-network activity that occurred when subjects actively engaged with the task. LC magnetization transfer contrast correlated with this state’s stability across experimental manipulations and with subjects’ propensity to enter into and remain in this state. These results provide new insight into LC’s role in driving spatiotemporal neural patterns associated with attention, and demonstrate that variation in LC integrity can explain individual differences in these patterns even in healthy young adults.

## Introduction

The locus coeruleus (LC) circuit is the main source of norepinephrine (NE) in the brain; it projects to the entire brain and is deeply involved in cognitive functions related to arousal including attention, stress, and the sleep-wake cycle (Aston-Jones & Cohen, 2005; X. Chen et al., 2014; Guedj et al., 2017; Langley et al., 2017; Sara, 2009; Song et al., 2017). For example, LC degradation, prevalent in normal aging, is thought to impair memory and cause cognitive reserve depreciation (Mather & Harley, 2016), and LC dysfunction is hypothesized to occur in prodromal stages of Alzheimer’s and Parkinson’s (Braak et al., 2003, 2011). In normal cognition, relationships among LC activity, arousal, attention, and performance have long been observed and are now classically characterized by the Yerkes-Dodson curve (Aston-Jones & Cohen, 2005): Moderate tonic LC firing rates correspond to optimal task performance while low and high tonic LC firing rates are associated with inadequate task performance because subjects are inattentive and distracted, respectively (Aston-Jones & Cohen, 2005).

But *how* do fluctuations in norepinephrine (due to fluctuations in LC activity) drive changes in brain states? Despite many observed correlations between LC activity and attention, the underlying mechanisms driving changes in network dynamics within this relationship are still not well understood (Aston-Jones & Cohen, 2005; Sara, 2009; Song et al., 2017). Characterizing these underlying mechanisms by studying normal cognition and LC engagement using computational models could provide foundational insight not only into healthy cognition, but also into disease states where LC structure and function is known to break down (Braak et al., 2003, 2011).

One primary measure of interest in developing such computational frameworks would be fluctuations in the LC itself, of course. This is highly challenging in awake, behaving humans in a noninvasive manner because of the extremely small size of the LC. The LC is only about 2 mm in diameter (**Figure 1**), meaning that even with recent advances in functional magnetic resonance imaging (fMRI) technology, LC blood oxygen level dependent (BOLD) signal can be incredibly noisy, as LC BOLD is especially susceptible to noise due to cardiac pulsation and respiration (Clewett et al., 2016; Glover et al., 2000; K. Y. Liu et al., 2017; Mather et al., 2017). Difficulties in accurately and precisely measuring the BOLD signal in LC render even sophisticated methods ineffectual. Seeking to quantify functional connectivity with LC, or using LC as a seed region for psychophysiological interactions analysis (O’Reilly et al., 2012), for example, might simply discover covariations with noise.

**Figure 1.**
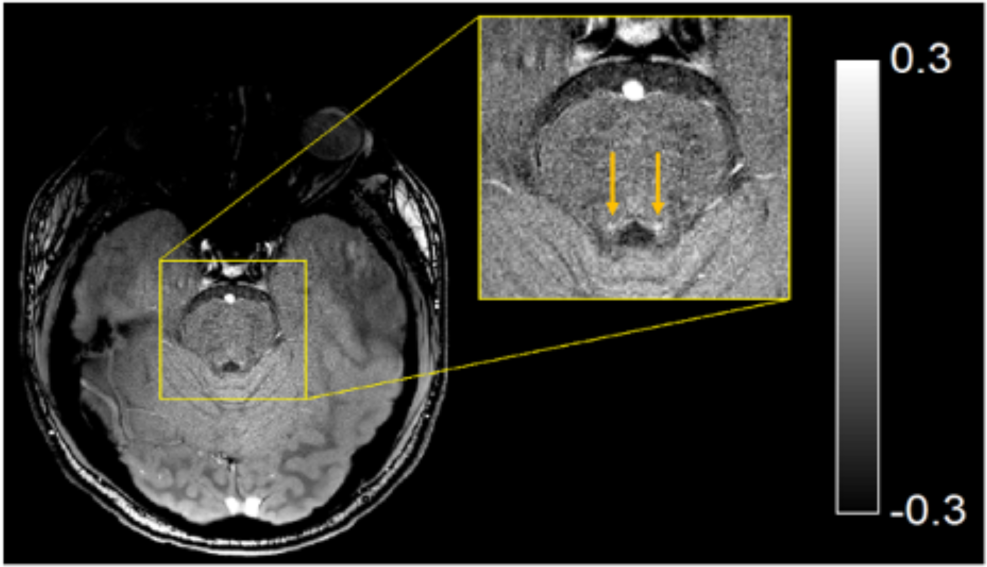
Neuroanatomical location of the locus coeruleus as seen in a magnetization transfer-prepared gradient echo image. The locus coeruleus (LC) is a 2mm diameter bilateral nucleus located in the brainstem, here illustrated by the two dots on either side of the region anterior to the fourth ventricle (X. Chen et al., 2014; Langley et al., 2017). The gray gradients used in the image illustrate magnetization transfer contrast (MTC), which is a unitless quantity. In general, voxels with negative values represent voxels with stronger magnetization transfer effects relative to the reference region. Shown here is a single subject in native space. The sequence was acquired with a MT-prepared gradient echo sequence described in the main text. The arrows indicate the position of LC.

To complicate matters more, there also exist individual differences in the neuronal density of LC even in young people (Keren et al., 2009). Although we are unaware of any documented relationships between memory performance and LC microstructure in younger adults (Langley et al., 2020, 2021; Mather & Harley, 2016) and neuronal loss in young adults is expected to be minimal if present at all (Manaye et al., 1995; Shibata et al., 2006; Zucca et al., 2006), variations in LC neuronal density may nevertheless lead to individual differences in the effectiveness (or effect size) of any manipulation designed to elicit fluctuations in LC activity, reducing group-level effect sizes further due to heteroscedasticity in post-manipulation behaviors across subjects. Indeed, there exist studies that have found relationships between LC integrity and memory performance in older but not younger adults, but where the between-groups interaction was not significant, which could be interpreted as indicating that the association might differ only in magnitude but not quality across groups if only sample sizes had been larger (Dahl et al., 2019).

One possible solution to these challenges is to use computational models that uncover and characterize repeating patterns in activity-based brain states, i.e. spatial patterns of co-activation that cycle repeatedly over a period of time. A popular tool for discovering these brain states is the hidden Markov model (HMM). HMMs can be used to discover patterns in complex, dynamic datasets, and are commonly used for weather prediction, computational biology, and finance (Eddy, 2004; Khiatani & Ghose, 2017; Zhang et al., 2019); recently, HMMs are becoming more popular in neuroimaging. Neuroimaging-based HMMs identify latent brain states which quantify network or nodal interaction as well as the probability of transitioning between those hidden states (Baker et al., 2014; S. Chen et al., 2016; Eavani et al., 2013; Lindquist et al., 2007; W. Liu et al., 2014; Ou et al., 2015; Robinson et al., 2010; Shappell et al., 2019; Stevner et al., 2019; Diego Vidaurre, Abeysuriya, et al., 2018; Diego Vidaurre, Hunt, et al., 2018; Diego Vidaurre et al., 2016, 2017). As a result, an HMM could be applied to fMRI data to identify the spatiotemporal characteristics of latent brain states as a function of LC up-regulation overall, rather than simply covariation with LC BOLD signal, in an attempt to characterize LC’s dynamic underlying relationship with arousal-modulated functions such as attention or sensitivity to stimulus salience.

Here, we modified a squeezing task, previously shown to induce sympathetic arousal and increase norepinephrine activity (Kozłowski et al., 1973; Lake et al., 1976; Mather et al., 2020; Nielsen et al., 2015; Nielsen & Mather, 2015; Vecht et al., 1978; Wallin et al., 1992, 1987) to create a pseudo-resting state fMRI paradigm to up-regulate LC (Hussain et al., 2019; Mather et al., 2020). We then used an HMM to characterize spatial patterns of co-activation among 31 ‘nodes’ comprising four known networks in the brain – the default mode network (DMN), dorsal attention network (DAN), fronto-parietal control network (FPCN), and salience network (SN) (Deshpande et al., 2011; Laird et al., 2005; Lancaster et al., 2007; Raichle, 2011) – as well as the LC itself (Langley et al., 2020, 2021). We then examined the states and their dynamic characteristics extracted via the HMM, including spatial patterns of brain activity and dynamics of state occupancy and transitions between states. We also used custom MRI sequences (Langley et al., 2017, 2020) to examine how individual differences in LC structure affect these measures.

The HMM was able to extract five stable brain states, including a highly stable state dominated by activity in the salience network that occurred when subjects engaged with the squeezing task; the stability of this state was correlated with LC neuronal density. Further, the propensity to dwell in this state, as well as the propensity to transition into this state from a state of relative deactivation, were both correlated with LC neuronal density. Together, our results reveal that our handgrip task was effective at changing the ease of transition into an arousal– or task-related state and the time spent in that state once achieved, and that individual differences in LC MTC explain important individual differences in the probability of attaining and maintaining neural activity states related to salience and attention, even in healthy young adults.

## 2. Methods

In this section, we first explain the experimental paradigm and neuroimaging data collection and analysis for both structural and functional data, including measures of LC neuronal density. We also introduce four attention-related networks and describe the HMM to be fitted. The next sections then describe how we approached answering questions about how activation of LC through the behavioral task affects brain network activity and state dynamics. This includes analyzing the HMM-derived brain states themselves as well as their evolution through time. We also performed exploratory pupillometry analyses on a subset of subjects for whom pupillometry data were useable (data loss due to data quality and acquisition issues led to an unfortunately large number of unusable datasets); these pupillometry analyses and results are presented in the **Supplementary Material.**

### 2.1 Datasets and Networks

#### 2.1.1 Experimental paradigm and neuroimaging & physiology data

##### 2.1.1.1 Participants

Thirty-one healthy adult human participants (18 females, mean age 25 years ± 4 years) enrolled in this study at the University of California, Riverside Center for Advanced Neuroimaging. Sample size was set to 30, which is within the range commonly used in studies of resting state dynamics (Deshpande et al., 2011), and to facilitate power analyses in a related study in the same subjects (not shown here) to inform future studies; one subject’s data could not be used due to technical difficulties during data acquisition, which led us to collect 31 subjects in total (30 with usable data). All subjects gave written informed consent and received monetary compensation for their participation, and had no history of neurological conditions or metal implants as required by inclusion criteria. All procedures were approved by the University of California, Riverside Institutional Review Board.

##### 2.1.1.2 Paradigm

The experimental paradigm is illustrated in **Figure 2A**. All subjects first underwent a five-minute pure resting state block (RS0) prior to any squeeze; this length has been shown to allow reliable measurement of resting state dynamics (Birn et al., 2013). Following this RS0 block, subjects underwent a 12.8-minute experiment where they alternated between resting state and bringing their dominant hand to their chest to squeeze a squeeze-ball at maximum grip strength (SQ1-RS5) (Hussain et al., 2019; Mather et al., 2020). This paradigm has previously been shown to reliably increase LC activity and facilitate attention to salient stimuli (Mather et al., 2020). All five squeeze periods lasted 18 seconds, while the interspersed five resting state periods had durations of two-, two-, five-, one-, and one-minute, respectively. SQ1 through RS5 occurred after the arousing stressor has been introduced, so we refer to them collectively as the post-arousal (PostAr) block. RS0 and PostAr blocks were collected separately within each condition for a total of four runs of functional data collection. In order to create a within-subject experimental design, all subjects underwent two sessions corresponding to two different conditions: one where they executed the squeeze (active condition) and one where they still brought their arm up to their chest but were instructed simply to hold the ball and not to squeeze it (sham condition). Condition order was pseudorandomly counterbalanced across subjects, and each condition occurred on a separate day. Following this resting state paradigm, subjects took part in an auditory oddball detection task, the details of which have been described elsewhere but which are not analyzed in the present project (Yaghoubi et al., 2019).

**Figure 2.**
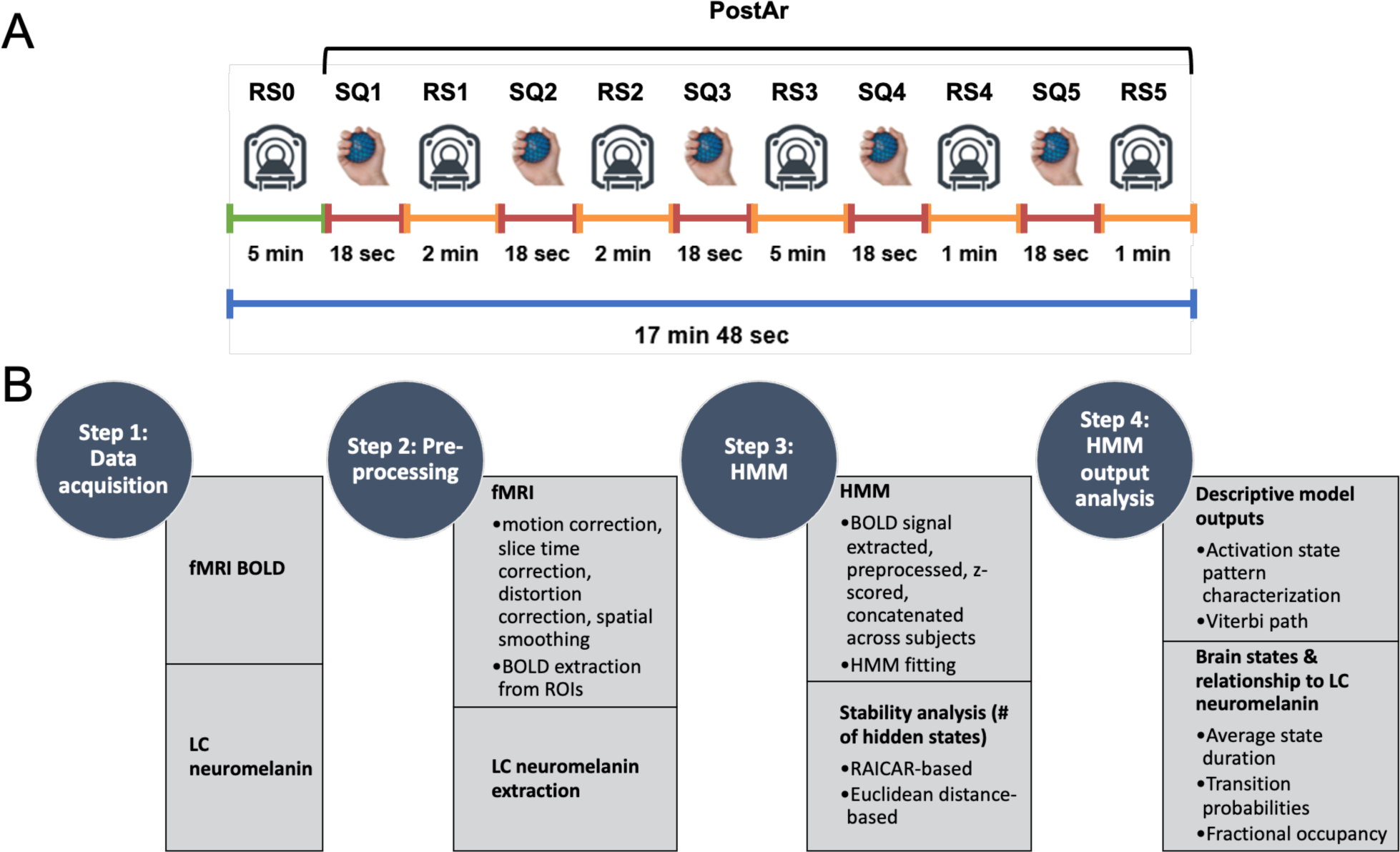
Experimental paradigm and data processing pipeline. (A) The experimental paradigm consisted of a pure resting state block and a longer alternating squeeze/resting state block to facilitate dynamic (time-varying) analysis (Nielsen & Mather, 2015). Subjects underwent two conditions: (1) an *active* squeeze condition, in which they lifted their arm to their chest and squeezed a squeeze-ball at maximum grip strength for the duration of each squeeze period; and (2) a *sham* control condition, in which they also lifted their arm to their chest during these periods but refrained from squeezing. Within the post-arousal (PostAr) block, RS refers to resting state periods, and SQ to periods in which subjects lifted their arm to their chest and either squeezed (active condition) or did not squeeze (sham condition). The RS0 blocks were used as a baseline for each condition. (B) The data acquisition and processing pipeline proceeded through four main steps, from data acquisition and pre-processing through HMM fitting and exploration of outputs. See main text for details of each step.

##### 2.1.1.3 Functional and structural neuroimaging data acquisition and preprocessing

Magnetic resonance imaging (MRI) data were collected on a Siemens 3T Prisma MRI scanner (Prisma, Siemens Healthineers, Malvern, PA) with a 64-channel receive-only head coil. fMRI data were collected using a 2D echo planar imaging sequence (echo time (TE) = 32 ms, repetition time (TR) = 2000 ms, flip angle = 77°, and voxel size = 2×2×3 mm^3^, slices=52) while pupillometry data (see **Methods**) were collected concurrently with a TrackPixx system (VPixx, Montreal, Canada). Anatomic images were collected using an MP-RAGE sequence (TE/TE/inversion time = 3.02/2600/800 ms, flip angle =8°, voxel size = 0.8×0.8×0.8 mm^3^) and used for registration from subject space to common space.

The functional data underwent a standard preprocessing pipeline in the functional magnetic resonance imaging of the brain software library (FSL): slice time correction, motion correction, susceptibility distortion correction, and spatial smoothing with a full width half maximum value of 2mm (Smith et al., 2004; Woolrich et al., 2009). Finally, all data were transformed from individual subject space to Montreal Neurological Institute (MNI) standard space using the following procedure in FSL (Smith et al., 2004; Woolrich et al., 2009). First, the T_1_-weighted image was skull stripped using the brain extraction tool. Next, brain-extracted T_1_-weighted images were aligned with the MNI brain-extracted image using an affine transformation. Finally, a nonlinear transformation (FNIRT) was used to generate a transformation from individual T_1_-weighted images to T_1_-weighted MNI common space (Smith et al., 2004; Woolrich et al., 2009). These steps correspond to steps 1 and 2 in **Figure 2B**.

##### 2.1.1.4 Regions of interest: attention-related networks

To examine the impact of LC up-regulation on network states and dynamics, we selected four networks often associated with resting state: default mode network (DMN), fronto-parietal control network (FPCN), dorsal attention network (DAN), and salience network (SN). DMN (a resting state network) and DAN (an attention network) were selected because squeezing ought to invoke a transition from the resting state into a task-positive state (Greicius & Menon, 2004); FPCN because it is linked to DAN and regulates perceptual attention (Dixon et al., 2018); and SN because it determines which stimuli in our environment are most deserving of attention (Mather et al., 2020; Menon & Uddin, 2010). Talairach coordinates for regions of interest (ROIs) within DMN, FPCN, and DAN were taken from Deshpande and colleagues (Deshpande et al., 2011) and converted to MNI coordinates while SN MNI coordinates were taken directly from Raichle’s 2011 paper (Deshpande et al., 2011; Laird et al., 2005; Lancaster et al., 2007; Raichle, 2011). Two ROIs from FPCN (dorsal anterior cingulate cortex and left dorsolateral prefrontal cortex) were excluded due to their close location to other ROIs. This step is shown in **Figure 2B**.

The fifth ‘network’ we included in fitting the HMM was the bilateral LC itself. LC was localized using the probabilistic atlas described in Langley et al. 2020 (Langley et al., 2017, 2020, 2021), not spheres as with the other networks. This atlas is specifically designed to allow comparable LC ROIs across subjects. Briefly, the locus coeruleus atlas was created by thresholding LC in MT-prepared GRE images and has been found to be reproducible in separate studies (Langley et al., 2020; Wengler et al., 2020). This atlas was segmented into rostral and caudal portions. Next, the entire LC atlas, rostral portion of the LC atlas, and caudal portion of the LC atlas were thresholded at a level of 0.5 and binarized. Because the MT images have contrast that is similar to those in T2-weighted images (i.e. white matter is dark and gray matter is bright), we elected to use a rigid-body transform with boundary-based cost function to derive a transform between MT and T1-weighted images. After the transform was derived, each registration was checked by visually assessing the overlap between each subject’s white matter mask and white matter regions in the MTC image. An example of this agreement is shown below (**Figure 3A**). We have previously shown that this procedure is reproducible and reduces mis-registration between T1-weighted and MT images (Langley et al., 2017; van der Pluijm et al., 2021; Wengler et al., 2020), and show here that our extracted BOLD signal for LC regions (both rostral and caudal) shows expected differences between active squeeze and sham control sessions (**Supplemental Material Section S2.2)**. This was implemented using FLIRT in FSL.

**Figure 3.**
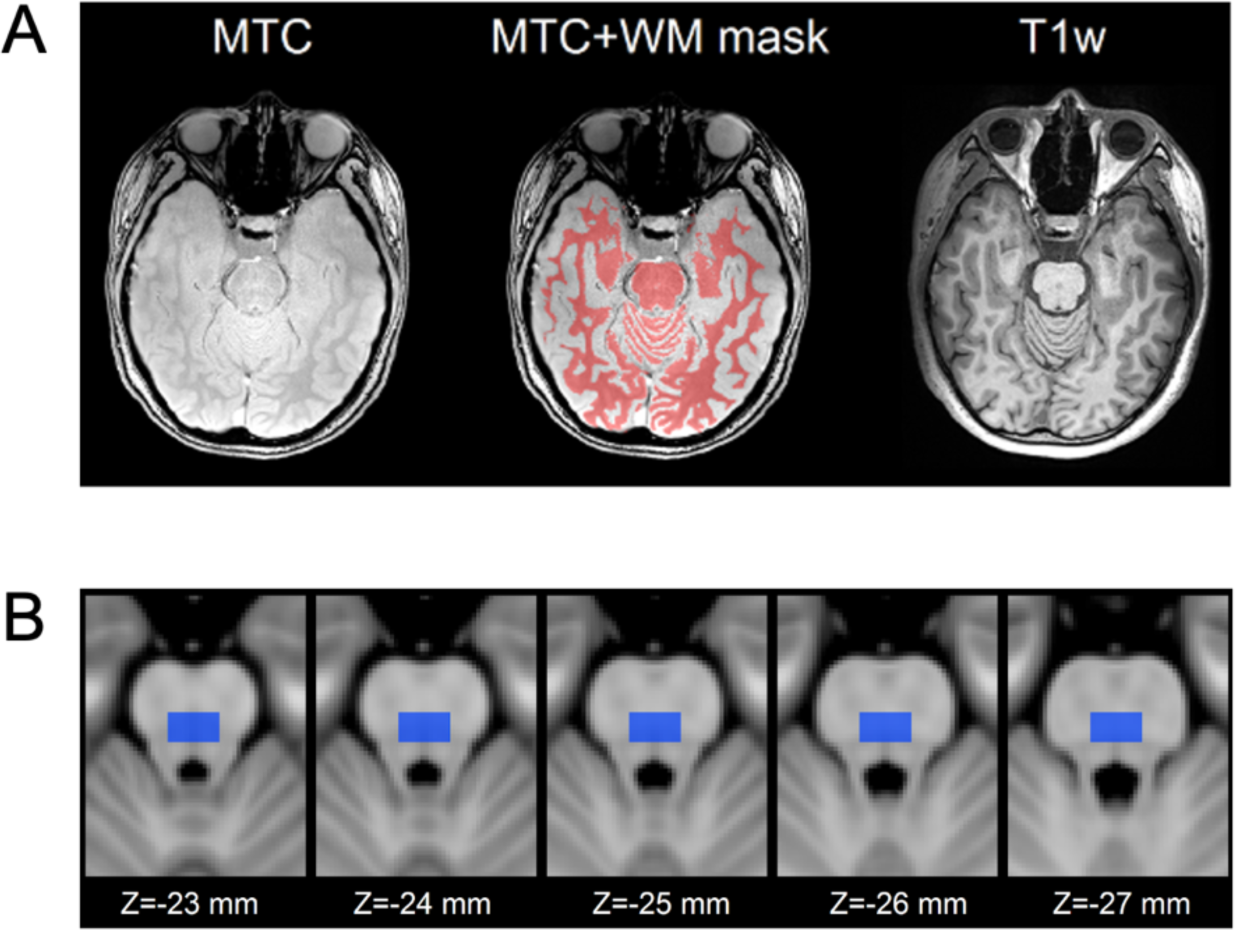
Demonstration of the LC imaging and localization process. (A) A comparison of magnetization transfer (MT) prepared gradient echo and T1-weighted images can be seen in the left and right columns, respectively. The center column shows white matter mask, segmented from the T1-weighted image displayed in red, overlaid on the MT prepared gradient echo image for a subject. There is good agreement between the white matter boundary mask and the white matter – gray matter boundary in the MT prepared gradient echo image, indicating accurate registration between T1-weighted and MT prepared gradient echo images. (B) The reference region used to calculate MTC in the MT images is shown in blue and overlaid on the 1mm T1-weighted MNI template. The reference ROI was defined as a 12 voxel by 6 voxel box in 5 slices anterior to the 4th ventricle on the 1mm T1-weighted MNI template.

**Table S1** shows the labels, and MNI coordinates for all networks and ROIs used in the analyses presented in the main text.

After preprocessing, all registrations (i.e. registrations between T1-weighted images and MNI space, fMRI images and T1-weighted images) were visually checked for each subject. In addition, the correspondence between the MNI T1-weighted template and fMRI images was also assessed. We detected no significant mis-registrations and the locus coeruleus atlas was not found to encroach in the 4th ventricle. Finally, for each subject, fMRI time series were extracted in each LC ROI.

##### 2.1.1.5 LC magnetization transfer data acquisition and preprocessing

Magnetization transfer (MT) prepared gradient echo images were used to compute LC magnetization transfer contrast (MTC) (reference **Figure 2B** for localization of this step in the processing and analytic pipeline). Data were acquired using a magnetization-prepared 2D gradient recalled echo (GRE) sequence: TE/TR = 3.10/354 ms, 416 × 512 imaging matrix, 162 × 200 mm^2^ (0.39 × 0.39 × 3 mm^3^) field of view, 15 slices, flip angle = 40°, four measurements, MTC preparation pulse (300°, 1.2 kHz off-resonance, 10 ms duration), and 470 Hz/pixel receiver bandwidth with a scan time of 10 minutes and 12 seconds (X. Chen et al., 2014; Langley et al., 2015). The four measurements were saved individually for offline registration and averaging. Slices on the MT-prepared GRE acquisition were prescribed perpendicular to the dorsal edge of the brainstem in the T_1_-weighted image. Two subjects chose not to participate in the MT-prepared gradient echo scans, so all MT-prepared GRE images analyzed in this project were done with n = 28.

To process the MT-prepared GRE data in FSL, images from the four GRE measurements were registered to the first image using a linear transformation tool in FLIRT and averaged (Smith et al., 2004; Woolrich et al., 2009). A transformation between this averaged MT-prepared GRE image and T_1_-weighted image was derived using a rigid body transform with boundary-based registration cost function in FLIRT. Prior to the rigid body transformation, the T_1_-weighted image was parceled into gray matter, white matter, and cerebral spinal fluid regions. The quality of each registration between T_1_-weighted and MT-prepared GRE images was assessed by overlaying the white matter-gray matter boundary from the T_1_-weighted image on the MT-prepared GRE image. No significant deviation was observed in all subjects. Contrast from the magnetization transfer preparation pulse, denoted MTC, was then calculated via

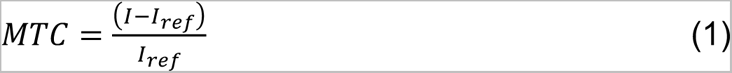

where *I* denotes the intensity of a voxel in the MT-prepared GRE image and *I*_ref_ refers to the mean intensity of a reference region in the NM-MRI image. To ensure consistent placement of reference region in MT-prepared GRE images across subjects, a reference region was drawn in the pons in MNI T_1_-weighted common space and then transformed to individual MT-prepared GRE images. This process ensures that the reference ROI has a similar location for all subjects and removes operator dependent bias in placement of the reference ROI and calculation of MTC. The reference region of interest (ROI) used in the analysis is shown in blue in **Figure 3B**. A LC atlas in MNI space was used in this study to localize the region around LC for MTC measurement (Langley et al., 2020). The LC atlas was transformed to MT-prepared GRE space using the aforementioned transformations and using a threshold level of 0.5. After binarizing, each subject’s mean MTC was measured in the LC ROI resulting in a total of n = 28 values. (Two subjects did not undergo the MT-prepared GRE scans.)

### 2.2 Hidden Markov model

#### 2.2.1 Model description and fitting procedures

To identify hidden brain state patterns and dynamics, and how they changed as a function of LC activity, we fitted a Gaussian hidden Markov model (HMM) to the functional neuroimaging dataset following previously-published methods (S. Chen et al., 2016; Hussain et al., 2022; Stevner et al., 2019; Diego Vidaurre et al., 2017). BOLD signals from the various ROIs (see **Methods Section 2.1.1.4**) were extracted, preprocessed, z-scored, and concatenated across subjects. Note that the data for RS0 and PostAr (SQ1-RS5) were z-scored separately within each condition because they formed four separate runs during acquisition, for a total of four z-scorings performed per subject: RS0 for active and sham, and SQ1-RS5 (PostAr) for active and sham. These fMRI time series were then concatenated timewise across all subjects to create a matrix of size (time * # subjects) x (# ROIs) and submitted as input to the hmmlearn python package to fit with standard procedures described elsewhere (Pedregosa et al., 2011). Briefly, the forward and Viterbi algorithms were used in conjunction to identify the most likely sequence of hidden states given the observable BOLD signal. The forward algorithm is used to calculate the ‘belief state’, i.e. the probability of a state at a given moment in time given the history of the evidence; the Viterbi algorithm is then used to estimate the most likely sequence of states, i.e. the most likely state *sequence* given the history of the observations. The Baum-Welch algorithm was then implemented to calculate the transition and emission probabilities of a given state (Jurafsky & Martin, 2009; L. Rabiner & Juang, 1986; L. R. Rabiner, 1989). The HMM was fitted on a group level to find the maximum number of global states that are visited by all subjects. Although this group-level analysis may ignore subject– or condition-specific states, the purpose here was to focus on shared states across subjects; further, fitting to individual subjects with a dataset this short can often result in unstable model fits or non-convergence. This step is shown in **Figure 2B** in the context of the rest of the processing pipeline.

#### 2.2.2 Number of hidden states

HMMs are fitted with an *a priori* defined number of hidden states, so we must first find that number before we can fit the model. For succinctness, in this section and the accompanying results section (see below) we refer to the number of hidden states of a particular HMM instantiation as the *model order.* To determine the optimal model order for the HMM used here, we employed two separate methods.

First, following previous approaches we adopted the Ranking and Averaging Independent Component Analysis by Reproducibility (RAICAR) method (S. Chen et al., 2016; Hussain et al., 2022; Z. Yang et al., 2008); this approach computes the ‘stability’ of recovered hidden state patterns to determine the appropriateness of a given *a priori* specified model order and therefore that model’s goodness of fit. Here, we examined the stability of model orders 3-15 (range based on previously established norms; (S. Chen et al., 2016; Hussain et al., 2022; Z. Yang et al., 2008)); As a second, confirmatory approach, we also employed Euclidean distances to assess model stability and model order. In this case, a smaller Euclidean distance between two matched states indicates a better correspondence between model initializations, i.e. a high goodness of fit. Further details of the RAICAR-based and Euclidean-based stability approaches are detailed in **Supplementary Material Section S2.1**.

### 2.3 Analysis of model outputs

We analyzed the outputs of the fitted HMM in a number of ways, summarized in step 4 of **Figure 2B**. First, we examined the state patterns themselves, i.e. how activity in each ROI changed as a function of state and across active versus sham control conditions. We also analyzed aspects of the trajectory through state space, including the Viterbi path itself, the percent of time spent in each state, the average state duration, and the transition probability matrices describing the model’s temporal evolution. These latter measures describing state trajectories were calculated separately for RS0 and the “post-arousal” (PostAr) block containing SQ1-RS5, because this latter portion of the scan occurred after the handgrip task (both active and sham). They were then compared across active versus sham conditions as well as compared to baseline dynamics during RS0. Below, we describe each of these approaches in detail.

#### 2.3.1 Exploring descriptive model outputs

##### 2.3.1.1 Activation state pattern characterization

The HMM fitting procedure directly outputs mean state patterns, i.e. the average activity shown by each ROI for a given hidden state recovered by the model. We also analyzed activation state patterns specific to the active and sham conditions. In accordance with Chen and colleagues’ (2016) method of state pattern acquisition, we acquired these spatial patterns by averaging the BOLD signal from TRs where the Viterbi path labeled a state to be active, for active and sham conditions separately. That is, for a given state within a given condition, we identified the TRs where a particular state was the most likely state to be occupied by the subject, and then averaged across all those TRs to produce an average BOLD signal in each ROI that can be tied to each state within that condition. We refer to this process as the *Viterbi averaging method*.

We compared the states recovered via the *Viterbi averaging method* across active versus sham conditions by first calculating the Pearson correlation coefficient between active and sham for each state, where state was defined as the average state found across all subjects. Because we cannot directly test for whether two states are the same, we rely on a lack of statistically significant difference to indicate insufficient evidence that the two states are dissimilar. We use Pearson correlations because we expect these relationships to be linear a priori.

Second, we again calculated the Pearson correlation coefficient between active and sham for each state (again because we assume these relationships to be linear), but this time within each subject so that we could perform statistical analyses. After Fisher-z transforming the correlation coefficients such that the distribution of data would not violate the normality assumptions of an ANOVA, we conducted a one-way repeated measures ANOVA on the transformed coefficients to seek a main effect of state. A significant main effect was followed by t-tests against 0 within each state.

Finally, we also calculated on an individual subjects basis the Euclidean distance between active and sham conditions for each state and each subject. The purpose of this analysis was to determine whether LC MTC correlates with how different the states are across this condition manipulation, so that these distances were then Spearman correlated with LC MTC across subjects; here we use Spearman correlations because there is no reason to expect the relationship between state distances and LC MTC to be linear a priori.

##### 2.3.1.2 Viterbi path

The Viterbi path – a direct HMM output – was used to qualitatively assess differences between active and sham conditions and to obtain qualitative insight into the temporal dynamics of the LC dataset. Since the active and sham conditions were fitted together, the model provided the hidden state sequence for concatenated active and sham conditions. We therefore separated the first and second halves of the outputted Viterbi path to examine the active and sham state sequences separately.

#### 2.3.2 Characterizing brain states and their relationship to LC structure

We next examined and characterized not only the states themselves, but how they behaved as a function of active versus sham conditions, and how these measures covaried with LC MTC across subjects. This allowed us to ask questions about how the squeezing task induced or changed transition into or occupancy of states related to attention or arousal (e.g., those characterized by dorsal attention or salience network activity), including how LC neuronal density (quantified by LC MTC) was related to these brain state characteristics.

We note again that each of the measures described in this section was computed separately for RS0 versus PostAr (SQ1-RS5; see **Methods Section 2.1.1.2**). This is because RS0 provides a baseline before any prompts are shown to either squeeze (active condition) or raise the hand to the chest (sham control condition), and so provide a benchmark against which we can look for *changes* in each measure, corrected for individual differences across subjects and conditions (and before application of the covariate of LC MTC). This approach also provides standardization across active and sham conditions, such that the critical questions centered on whether each metric’s deviation from baseline is dependent on whether the subject was actively squeezing or not.

For all metrics described in this section, the first omnibus test typically took the form of computing the relevant metric separately in the active and sham conditions and in the RS0 and PostAr blocks, such that every subject had four measures of the metric in question. To do baseline correction within each scanning session (which occurred on two separate days; see **Methods Section 2.1.1.2**) we then subtract the RS0 metric from the PostAr metric, separately in the active and sham conditions. The resulting difference score is submitted to statistical tests as appropriate, detailed in the following sections.

Finally, to determine the degree to which LC structure could help characterize the relationships found via the analyses described below, we calculated the Spearman correlation between LC MTC and these measures; as above, Spearman correlations were chosen because we had no *a priori* reason to believe LC MTC should be linearly related to any of these measures.

Below we describe these analyses in more detail.

##### 2.3.2.1 Average state duration

After the hidden states were extracted and characterized, we were interested to know whether the propensity to dwell in each state, once reached, changes across conditions or states. Thus, we defined average state duration as the mean time spent in a state once a subject entered that state. This was computed for each subject by calculating the number of consecutive TRs spent in a state once the subjects entered it, and then averaging this number across the total number of times the subject entered that state, separately for the active and sham conditions and also separately within RS0 and PostAr blocks. This leaves us with four average state duration measures for each subject: Dur_RS0,active_, Dur_RS0,sham_, Dur_PostAr,active_, and Dur_PostAr,sham_. Finally, we subtracted RS0 from PostAr within each of the active and sham conditions to get a difference score, i.e.

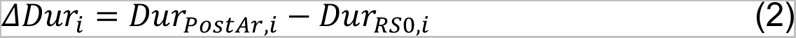

with *i* referring to condition, either active or sham. The resulting difference scores Δ*Dur* were submitted to a 2 (condition: active vs sham) x 5 (state) repeated measures ANOVA. We then conducted step-down ANOVAs and t-tests as appropriate; Greenhouse-Geisser corrections for sphericity violations were used as needed.

Finally, to examine the degree to which changes in average state duration as a function of condition were related to LC MTC across subjects, we computed the Spearman correlation between LC MTC and average state duration within subsets of the data (condition and state) as appropriate.

##### 2.3.2.2 Transition probabilities

A second question concerns how state transition characteristics change as a function of active versus sham condition, and the degree to which these changes are related to LC MTC. A *transition probability matrix* (TPM) provides a summary of all the dynamics observed in the Viterbi path, in that the value contained in *TPM*_*ij*_ describes the probability of switching from state *i* to state *j* at any given moment in time. The TPM directly outputted from the HMM fitting procedure provides a general overview of transitions for both conditions as a whole, and also includes baseline (RS0) effects since it reflects the fitting procedure applied to all data concatenated across subjects and conditions. However, we were interested in changes in transition probabilities across the several manipulations of our task.

First, we were interested to see how the TPM changed from baseline once the handgrip task was introduced, i.e., during PostAr for both active and sham conditions. To calculate this, we concatenated Viterbi paths within each of the active and sham conditions, respectively, during the PostAr block and identified the number of times a subject transitioned out of a certain state and into another, then divided that value by the total number of transitions in the block (374). To remove the effects of baseline, the TPM was calculated similarly for RS0 separately in each condition, and then subtracted from the overall PostAr transition probability matrix (again separately in each condition) to produce a *relative transition probability matrix* (RTPM) for each condition *c*∊{*Active, Sham*}:

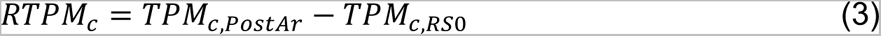

which describes the transition probabilities in the PostAr block relative to whatever baseline was set during RS0 for that subject during that scanning session.

Next, we checked for differences between RTPM_Active_ and RTPM_Sham_. To do this, we first computed the Euclidean distance between RTPM_Active_ and RTPM_Sham_ for each subject via

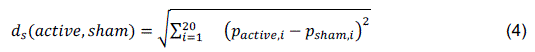

where *d*_s_ is the Euclidean distance between the transition probabilities for a particular transition *i* ∊ {*S*1 → *S*2, *S*1 → *S*3, …, *S*5 → *S*4} excluding self-transitions (i.e., 20 pairwise transitions in total), *p* is the transition probability for that particular state transition, and *s* ∊ {*subjects*}. *d*_s_ thus indexes the total distance between the active and sham RTPMs for a particular subject *s*, across all pairwise transitions. We compared the distribution of these distances to 0 using a nonparametric Wilcoxon sign rank test; this process is akin to a nonparametric paired t-test (because distances are by definition positive and so are not normally distributed).

We were also interested in which particular pairwise transitions might drive these differences, so we next computed *p_active,i_* − *p_sham,j_* for all pairwise transitions *i* → *j* within RTPM_Active_ and RTPM_Sham_ and for each subject, and tested whether each of these distributions was different from 0 across subjects with a series of Wilcoxon sign rank tests (again, parametric tests are inappropriate due to a priori violations of normality assumptions, as these are differences in probabilities which range between 0 and 1).

Finally, to determine whether LC MTC was related to changes in pairwise transition characteristics – that is, differences between RTPM_Active_ and RTPM_Sham_ – we Spearman correlated the differences between RTPM_Active_ and RTPM_Sham_ with LC MTC across subjects.

##### 2.3.2.3 Fractional occupancy

Finally, we were also interested in the percent of overall time that subjects spent in each hidden state. Was one state dominant over another, and did this change as a function of active versus sham condition and/or was it correlated with LC MTC? This is especially important because any differences in transition probabilities or state duration must be interpreted with respect to how much time, overall, a subject spent in each state: If a particular state becomes highly dominant in one condition over another, that might also carry increases in duration or transition probabilities into that particular state.

Therefore, we defined *fractional occupancy* (FO) as the proportion of time spent visiting a state. Fractional occupancy was calculated by first counting the total number of TRs each subject spent in a certain state for each block, RS0 and PostAr; that value was then divided by the total number of TRs in each block (150 for RS0 and 375 for PostAr) to obtain the fractional occupancy of each state separately within either RS0 or PostAr. We performed this calculation separately in the active versus sham conditions, and then subtracted RS0 from PostAr as done previously for average state duration. That is, we computed

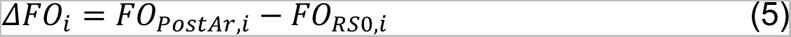

with *i* again referring to condition, either active or sham. Following the previous design for average state duration, the resulting difference scores Δ*FO* were submitted to a 2 (condition: active vs sham) x 5 (state) repeated measures ANOVA. We then conducted step-down ANOVAs and t-tests as appropriate; Greenhouse-Geisser corrections for sphericity violations were used as needed, and any outliers more than 3 standard deviations from the mean in any condition were discarded. Finally, to examine the degree to which changes in fractional occupancy as a function of condition were related to LC MTC, we again computed the Spearman correlation between LC MTC and fractional occupancy within subsets of the data (condition and state) as appropriate.

## 3. Results

### 3.1 Establishing foundations

#### 3.1.1 LC structural integrity results

We computed LC magnetization transfer contrast (MTC) using MT-prepared GRE images as a surrogate marker for LC neuronal density.Across subjects, the mean (± s.d.) LC MTC was 0.1212

± 0.0220. These values were correlated with the various model outputs as described in **Methods**.

We then confirmed that the correlation between rostral and caudal regions of LC was significant to ensure that the LC localization procedures were successful. These values are shown in **Table 1**.

**Table 1.** Correlation between rostral and caudal LC regions as a function of block and session. * p < 0.05. All significant effects survive correction for multiple comparisons via the Benjamini-Hochberg method.

|  | **Active** |  | **Sham** |  |
| --- | --- | --- | --- | --- |
|  | **Pearson's r** | **p** | **Pearson's r** | **p** |
| **RS0** | 0.5495 | <.001* | 0.5189 | <.001* |
| **PostAr** | 0.5969 | <.001* | 0.5954 | <.001* |

**Table 2.** Group-defined states show differences between active and sham conditions, quantified by the lack of statistically significant Pearson correlation coefficients (r) between active and sham versions of each state except for two states: S1, the DMN-dominant state (although this would not survive correction for multiple comparisons via the Benjamini-Hochberg method), and S4, the SN-dominant state, showed significant correlations between active and sham. * p < 0.05.

|  | Pearson's r | p |
| --- | --- | --- |
| **State 1** | 0.3763 | 0.0369* |
| **State 2** | -0.1738 | 0.3499 |
| **State 3** | 0.2447 | 0.1846 |
| **State 4** | 0.7887 | < 0.001* |
| **State 5** | 0.1739 | 0.3494 |

#### 3.1.2 Determination of number of hidden states

**Figure 4** shows the results from performing the RAICAR-based and Euclidean distance-based stability analyses (**Methods Section 2.2.2: Supplementary Material Section 2.1**) for model orders (number of hidden states) 3-15 using three initializations. Both plots indicate that five states were best for this investigation and produced highly stable state discovery, indicating high goodness of fit. This was the maximum model order where the stability values for all states remained above the predetermined threshold, and where the Euclidean distance remained as low as possible (zero) before dramatically increasing. Chen et al. 2016 (S. Chen et al., 2016) and Yang et al. 2008 (Z. Yang et al., 2008) both used the RAICAR-based method, explored 236 ROIs and 162 independent components respectively, and employed a stability threshold of 0.8 (S. Chen et al., 2016; Z. Yang et al., 2008). As discussed above we used a 0.9 threshold because we examined substantially fewer ROIs so our analyses and interpretations could afford to be more stringent. However, whether we used a threshold of 0.9 or 0.8 at least one stability value for model orders six and above fell below 0.8; so, a 5-state model was selected via both standards.

**Figure 4.**
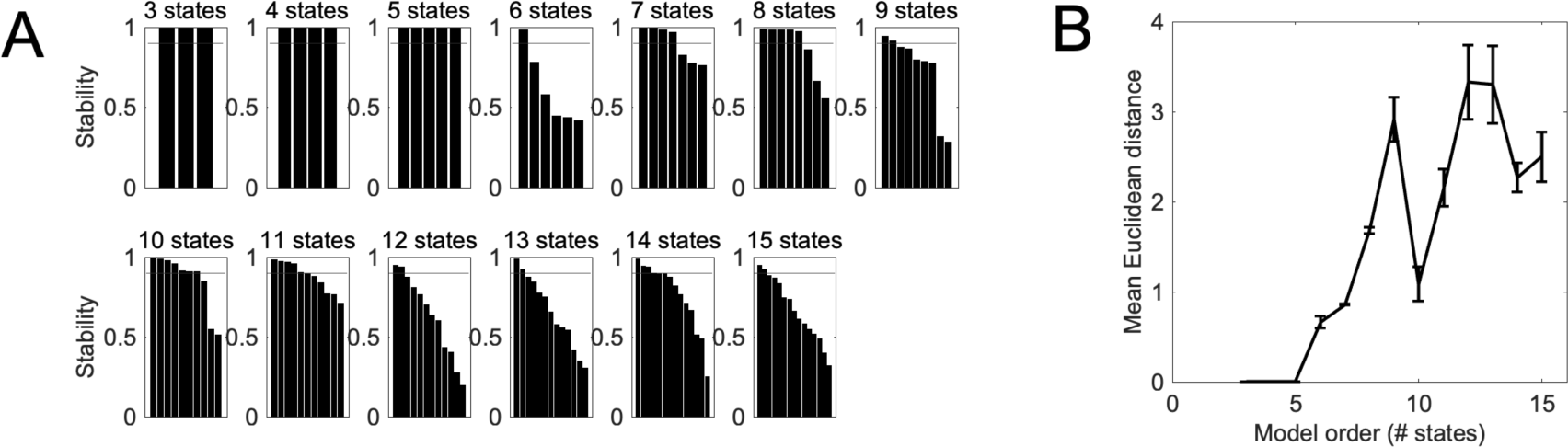
Results of stability analysis to determine HMM model order (number of hidden states). Stability analysis results via both the (A) RAICAR-based (B) and ED-based methods for model orders 3-15 indicate that a 5-state model was best, as this is the largest model order where the stability values were above the 0.9 threshold (thin horizontal lines) in the RAICAR-based results and the mean Euclidean distance was as small as possible with the largest number of states in the Euclidean distance-based results.

These methods determined that a 5-state model fit the data best (i.e., the discovered states were stable and demonstrated the best possible goodness of fit), and so all subsequent analyses and results presented here are for a 5-state HMM.

### 3.2 Descriptive model outputs

#### 3.2.1 Activation state pattern characterization

**Figure 5A** shows the activation state patterns recovered by the fitted HMM. State 1 (S1) appears to represent a DMN-dominant state because DMN showed the highest level of activation, while State 2 (S2) corresponds to an attention-dominant state since DAN and SN showed the highest levels of activation. State 3 (S3) shows all networks investigated to be activated, and State 4 (S4) can be labeled as the squeeze/sham state because it was prevalent during the squeeze/sham periods of the paradigm (**Figure 6**). Because this state was also prevalent during the “squeeze” periods of the sham condition, where the subjects lifted their arm to their chest but refrained from squeezing, S4 will be referred to as the task-related state. Reflecting that S4 is related to the task (which is designed to induce arousal), S4 also shows a dramatic increase in the activation level of some ROIs in the attention networks, DAN and especially SN – specifically the right anterior prefrontal cortex and left insula – as well as relative *deactivation* of LC. Finally, all networks were deactivated in State 5 (S5), perhaps because another network not examined in this investigation was activated (e.g., visual networks). Furthermore, we note that many of these states showed qualitative relationships to one another. For example, S1 and S4 visually appear to have similar patterns, but with opposite signs, such that DMN showed high levels of activation in S1 but was deactivated in S4. Conversely, right anterior prefrontal cortex and left insula in SN were deactivated in S1 but were activated in S4.

**Figure 5.**
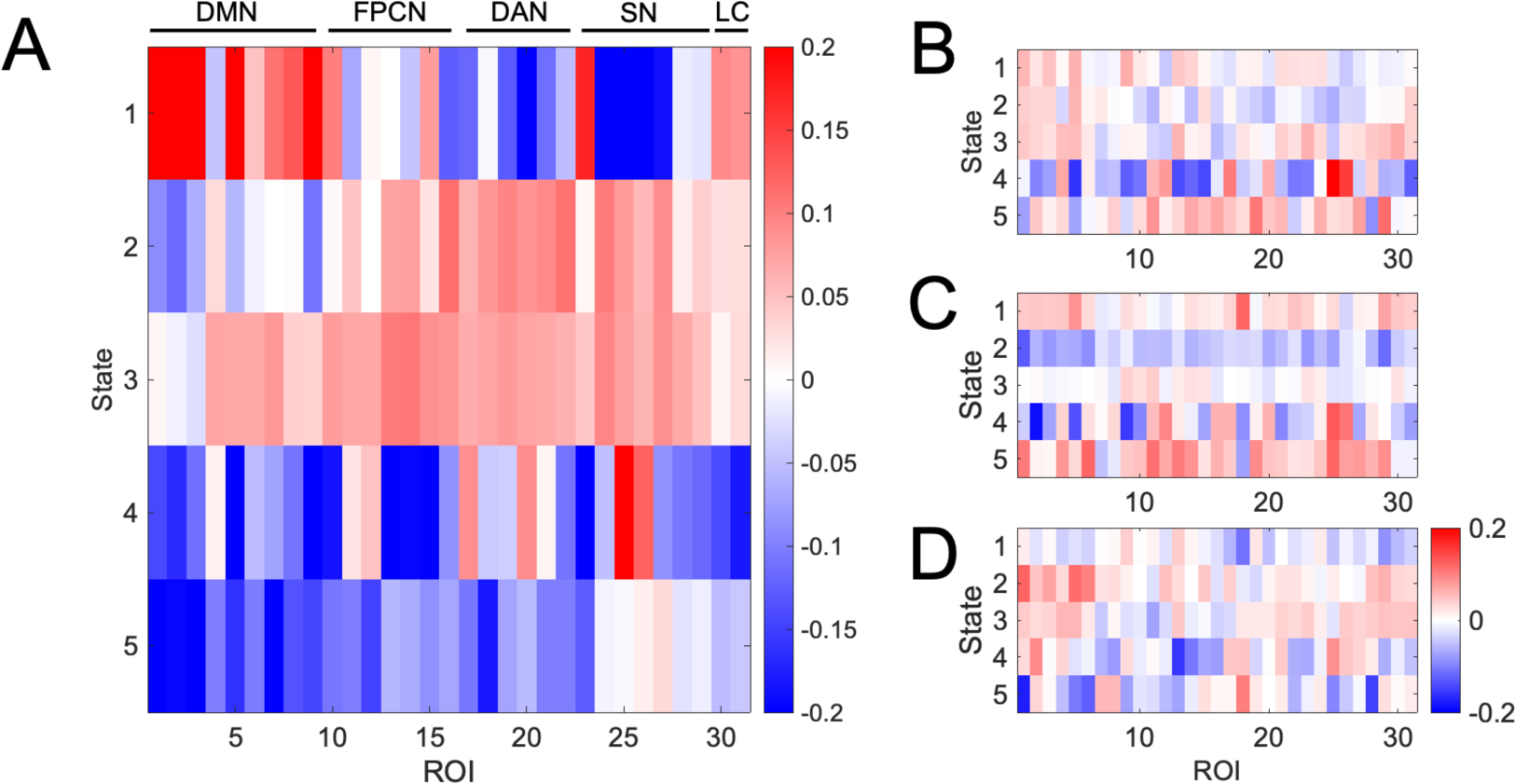
The five stable activation state patterns extracted by the fitted HMM. **(A)** Activation state patterns show interpretable qualitative patterns, such that state 1 (S1) appears to be DMN-dominant, S2 appears SN– and DAN-dominant, S3 shows broad activation across all networks, S4 appears to reflect a state dominated by sharp increase in SN activity (and some DAN engagement) with relative deactivation in all other networks, and S5 shows broad deactivation across networks. **(B)** and **(C)** show activation state patterns as a function of active versus sham condition, respectively, found by the *Viterbi averaging method* (see **Methods**). Activation state patterns for **(B)** active and **(C)** sham conditions show some qualitative and quantitative similarities (especially in S4), but also display important differences, shown in **(D)** which displays active minus sham. Units are normalized BOLD. See main text for detailed analysis.

**Figure 6.**
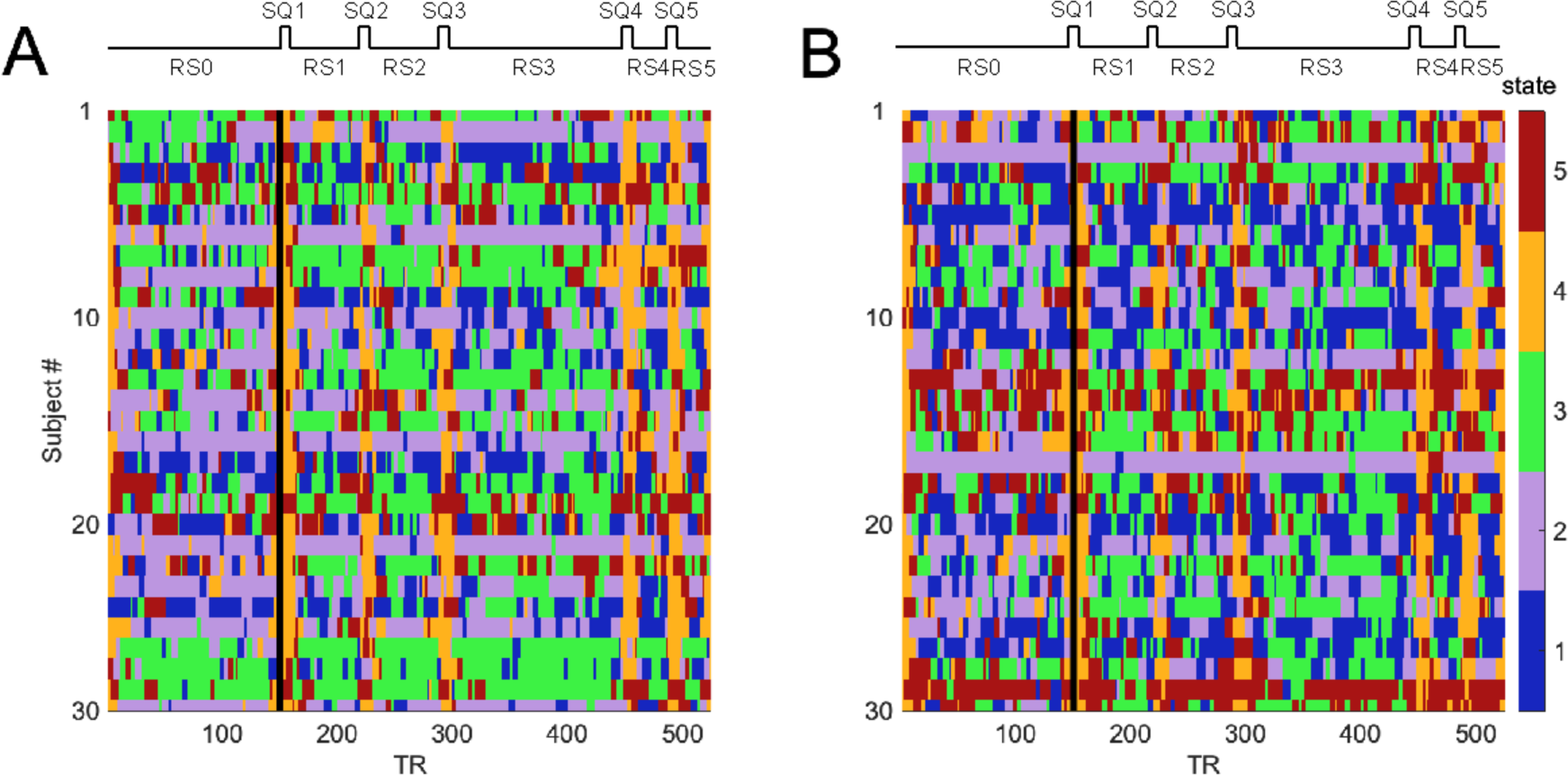
Viterbi paths for active and sham conditions. Visually, there appear to be meaningful differences in the state trajectories between **(A)** active and **(B)** sham conditions; these differences are quantitatively tested in the next sections. Vertical black lines indicate the break between RS0 (before any squeezing or hand-raising) and PostAr (after the first squeeze/hand raise) blocks (see **Methods Section 2.1.1.2**).

States 1 and 4, the DMN-dominant state and the SN-dominant state, respectively, show quantitative similarities between conditions, but the other states do not (**Figure 5D**). At the individual level, we repeated this process and correlated active and sham for each subject and each state. All but S4 showed a lack of significant correlation between active and sham conditions. This was revealed by a one-way repeated measures ANOVA with state as a factor, testing the Fisher-z transformed correlation coefficients between active and sham. This ANOVA showed a main effect of state (F(4,108) = 5.803, p < 0.001). We followed this with step-down t-tests against 0 to explore which state(s) were significantly correlated across active versus sham condition (**Table 3**).

**Table 3.** Mean Pearson correlation coefficient (r) and t-test results for comparing states between active and sham conditions on a subject-by-subject basis. S4 showed a high degree of stability (significantly positive Pearson correlation coefficients) between the active and sham condition. All statistical tests were performed after the correlation coefficients were Fisher z transformed, but the means and standard deviations are presented in raw correlation coefficient units. * p < 0.05. The significant test (State 4) survives correction for multiple comparisons via the Benjamini-Hotchberg method.

| | Mean<br>Pearson's r | $\sigma$ | t | p |
| --- | --- | --- | --- | --- |
| **State 1** | 0.1770 | 0.3436 | 0.1888 | 0.8594 |
| **State 2** | 0.0086 | 0.3100 | 0.0584 | 0.9571 |
| **State 3** | -0.0026 | 0.2998 | 0.7567 | 0.4915 |
| **State 4** | 0.2659 | 0.3360 | 5.4652 | 0.0055* |
| **State 5** | 0.0113 | 0.3143 | -0.7775 | 0.4803 |

Of the five t-tests against 0, only S4, the SN-dominant state, reached significance. These tests demonstrate that there are likely significant variations in the state patterns between active versus sham conditions (i.e., a lack of evidence for significantly *similar* patterns), but S4 remained highly stable across this manipulation even at the individual subject level. Although this may seem surprising given that the experimental manipulation was at its ‘largest’ during the squeeze periods (subjects either squeezed or held the ball to their chest), further scrutiny suggests that the downstream effects on the other networks are expected to exhibit the most variation as a function of the experimental squeeze manipulation. That is, because of the difficulties in measuring LC activity or connectivity directly, we should expect to see differences in states as a function of condition not precisely at the point of LC up-regulation, but in downstream effects on network activation patterns throughout the subsequent resting-state periods of the scan. Thus, going forward, we can see that S4 is indeed a stable state (occurring during the task) that makes a strong anchor point to focus the remaining analyses.

We can also characterize the differences between active and sham in the state patterns themselves. S1 (DMN-dominant) displays higher activity levels in the sham than in the active condition, while the opposite is true for S2 (attention-dominant). That S2 has higher activity in the active than sham condition suggests that LC activation in the active condition may have increased attention and/or arousal; this is also consistent with the observation that S3 (all networks activated) shows higher activity in active than sham, while the opposite is true for S5 (all networks deactivated). And even visually, one can see that the smallest differences between active and sham appear to occur in S4, consistent with the statistical analysis above.

Finally, we calculated the Euclidean distance between active and sham conditions for each state and each subject, and then correlated these distances with LC MTC to determine whether LC MTC could explain the effects of the active squeeze manipulation. We found that only S4 showed a trending relationship with LC MTC (**Table 4**), but importantly that this trending relationship was an inverse relationship: higher LC MTC was associated with smaller distance (i.e., greater degree of similarity) between the active and sham condition instantiations of this state. This finding suggests that LC MTC may be related to a capacity to stabilize this state of SN dominance, and that individual differences in LC MTC even in younger adults are related to significant differences in states of neural activity that are particularly responsive to stimulus salience.

**Table 4.** Spearman correlation coefficients between LC MTC and the Euclidean distance separating state activation patterns during active and sham conditions as a function of state. S4 showed a trending anti-correlation between LC MTC and distance, i.e. that higher LC MTC was associated with smaller distance between active and sham conditions for the activity pattern associated with this state. ^†^ p < 0.1

|  | Spearman's r | p |
| --- | --- | --- |
| **State 1** | -0.0482 | 0.8075 |
| **State 2** | -0.0757 | 0.7066 |
| **State 3** | 0.0077 | 0.9700 |
| **State 4** | -0.3629 | 0.0584<sup>†</sup> |
| **State 5** | 0.1910 | 0.3287 |

**Table 5.**
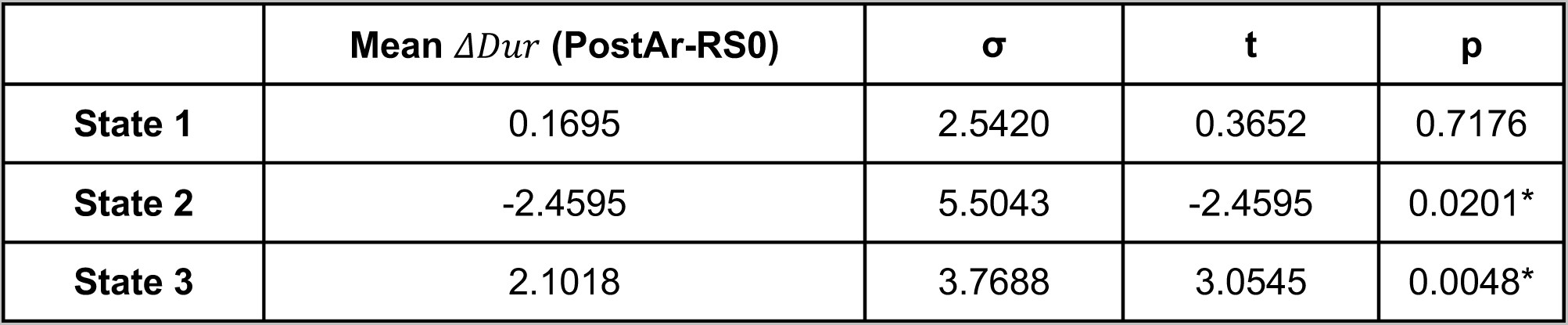
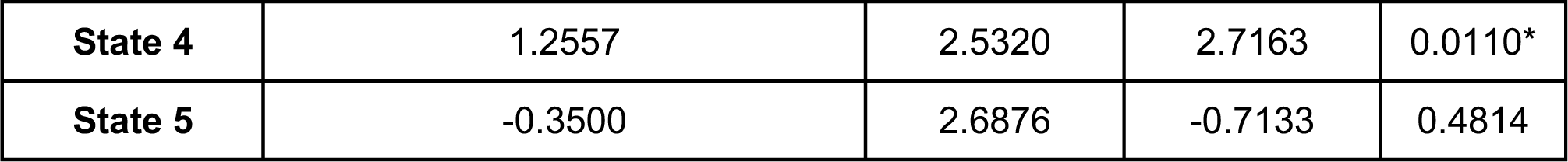
Average changes in durations and t-test results for comparing state duration changes relative to baseline across state, collapsed across conditions. Three states (S2, S3, and S4) showed significantly different durations in PostAr relative to baseline (RS0): S2 showed significantly shorter durations in PostAr, while S3 and S4 showed significantly longer durations in PostAr. See main text for details. * p < 0.05. All significant effects survive correction for multiple comparisons via the Benjamini-Hochberg method.

| | Mean $\Delta Dur$ (PostAr-RS0) | $\sigma$ | t | p |
| --- | --- | --- | --- | --- |
| State 1 | 0.1695 | 2.5420 | 0.3652 | 0.7176 |
| State 2 | -2.4595 | 5.5043 | -2.4595 | 0.0201* |
| State 3 | 2.1018 | 3.7688 | 3.0545 | 0.0048* |
| State 4 | 1.2557 | 2.5320 | 2.7163 | 0.0110* |
| State 5 | -0.3500 | 2.6876 | -0.7133 | 0.4814 |

#### 3.2.2 Viterbi path

The Viterbi paths for active and sham conditions are illustrated in **Figure 6**. Remarkably, the squeeze periods within the *post-arousal* (PostAr) block (after the first squeeze or hand-raise had occurred; see **Methods Section 2.1.1.2**) were obvious, as almost all subjects visit S4 (the orange state) in both active and sham conditions for almost the entire length of the PostAr blocks at time points 151-159, 220-228, 289-297, 448-456, and 487-495 – visible as vertical orange bars running through almost all subjects.

Visually, we see hints that the Viterbi paths may be different between active and sham conditions. For example, S3 seems more prominent in active than sham, whereas S1 may be more prominent in sham. S5 also appears to occur more often in sham. Rather than rely on visual inspection, we quantitatively compared the evolution and dynamics of the active and sham Viterbi paths and describe the results in the next sections.

### 3.3 Brain state characteristics & relationship to LC structure

We next examined several dynamic aspects of the state space trajectories themselves. These measures were also correlated with LC MTC. See **Methods** for details.

#### 3.3.1 Average state duration

The first component we examined were the baseline-corrected average state durations (see **Methods Section 2.3.2.1**). An omnibus 2 (condition: active vs sham) x 5 (state) ANOVA showed a main effect of state (F(4,116) = 11.155, p < .001) but no main effect of condition (F(1,116) = .355, p = .556) and no interaction (F(4,116) = .231, p = .921) (**Figure 7A**). This pattern suggests that once subjects entered a state, the amount of time they dwelled in that state differed from one state to the next but did not differ across the active versus sham manipulation. We therefore collapsed across conditions and recalculated average state duration on the collapsed dataset, and followed this with a one-way repeated measures ANOVA testing for a main effect of state on duration.

**Figure 7.**
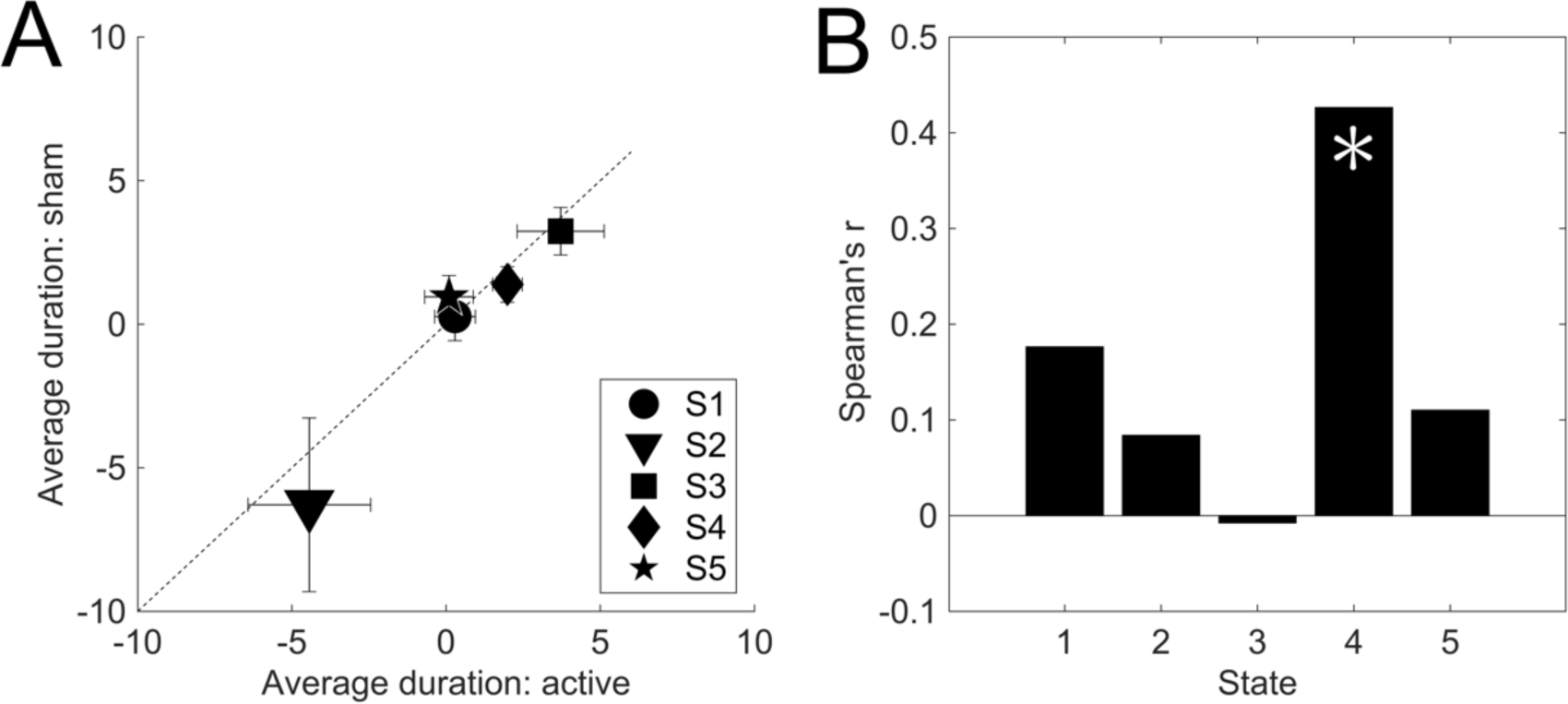
Average baseline-corrected state durations (Δ*Dur*) in the active and sham conditions, and correlations with LC MTC across subjects. **(A)** The 2 (condition: active vs sham) x 5 (state) repeated measures ANOVA revealed no main effect of condition but a main effect of state, which is visible in the nearness of the points to the identity line. **(B)** Average durations for S4, the SN-dominant state, showed significant (p = 0.0246) correlation with LC MTC: the more neurons present in a subject’s LC, the longer the participant tended to dwell in S4 (after correcting for baseline). No other correlations with LC MTC reached significance. Error bars in **(A)** show the standard error of the mean. See main text for details. * p < 0.05

This process revealed a main effect of state (F(4,116) = 7.236, p < .001), which we then followed with five t-tests against 0 to see which states had higher or lower average duration in the PostAr block relative to baseline (RS0, before any squeezing or hand-raising had occurred; see **Methods Section 2.1.1.2**). These tests revealed that three states were on average significantly longer or shorter in duration in PostAr than in RS0 (**Table 6**): S3 (“everything is activated”) and S4 (SN-dominant task state) were significantly longer in PostAr than in RS0, while S2 (attention-dominant state) was – perhaps surprisingly – significantly shorter in PostAr than in RS0. S2 being longer in RS0 than PostAr may make sense because RS0 occurred at the beginning of a relatively long scan session, so subjects may have been alerted by the sounds of the scanner or instructions from the experimenter. S3 being longer in PostAr may similarly be related to the short duration of RS0, as well as the relatively less time spent in this state (see below). S4 being longer in PostAr makes sense also because this is where the squeeze periods occurred, and so S4 rarely occurred during RS0 and only at the very beginning of this first scan run (**Figure 5**) and therefore its duration was truncated.

**Table 6.** Spearman correlation coefficients (r) comparing LC MTC to average state duration change from baseline, collapsed across conditions. The change in duration (relative to baseline) of S4, once entered into, was significantly correlated with LC MTC across subjects. See main text for details. * p < 0.05. However, we note that the only significant effect, S4, would not survive correction for multiple comparisons via the Benjamini-Hochberg method.

| | Spearman's $r$ | $p$ |
| --- | --- | --- |
| **State 1** | 0.1762 | 0.3680 |
| **State 2** | 0.0837 | 0.6708 |
| **State 3** | -0.0077 | 0.9700 |
| **State 4** | 0.4264 | 0.0246* |
| **State 5** | 0.1100 | 0.5773 |

Finally, we also wanted to see whether LC MTC could predict changes in state duration from baseline, across subjects. To answer this question, LC MTC was correlated with the average duration of all five states after (**Figure 7B**) collapsing the data across conditions as above since this factor was nonsignificant in the omnibus ANOVA. The Spearman’s r values in **Figure 7B** were acquired by correlating LC MTC with the difference in average duration across baseline, i.e. *corr*(Δ*Dur, LC MTC*) (concatenated across conditions).

Here, we observed only one significant correlation between LC MTC and the average duration of S4 (SN-dominant state) (**Table 6**). This is particularly noteworthy as S4 occurs in both active and sham conditions as a direct consequence of the task manipulation, and reflects engagement with the motor task. That is, this result shows that the integrity of a subject’s LC impacts how long that subject persists in the SN-dominant state, regardless of the active versus sham condition manipulation: the higher the inferred neuronal density in a given subject’s LC, the longer they persisted in an SN-dominant state once they achieve it. We noted in our first characterization of S4, above, that S4 contains targeted increases in activity in two nodes in particular, the right anterior prefrontal cortex and left insula, and relative deactivation of LC. Nevertheless, we can see that LC structure significantly predicts how long the aroused state persists once it is achieved. As we will see in the next sections, this focus on S4 as being especially sensitive to task demands and LC MTC is maintained through the next analyses.

#### 3.3.2 Transition probabilities

##### 3.3.2.1 General results for transition probabilities

Next, we asked whether transition characteristics between pairs of states were modulated by our active versus sham condition manipulation, and how these changes might be related to LC MTC. This analysis involves computing the transition probability matrices (TPMs), relative transition probability matrices (RTPMs), and their differences as a function of condition (see **Methods Section 2.3.2.2** for details).

First, we can visually examine the average TPMs from RS0 and PostAr, in the active and sham conditions (**Figure 8A-D**). Visually, we can see reasonable covariation among all of these, suggesting that the pairwise transitions are relatively stable across all blocks and conditions. Recall, however, that RS0 occurred on different days for both active and sham conditions. Therefore, to remove baseline effects, we really want to examine the RTPMs, not the TPMs. RTPMs are defined as the difference in TPM between PostAr and RS0, i.e. how much the TPM changes as a result of the introduction of the squeezing (active) or hand-raising (sham) task relative to baseline transition probabilities occurring during the first resting state period (RS0). In this case, baseline was important to consider because one subject (or even one condition’s RS0 in one subject, given that active and sham were collected on different days; see **Methods Section 2.1.1.2**) may intrinsically have had a higher switching rate than another subject or condition’s RS0. To understand how the handgrip task affected the transition probabilities, we thus removed any baseline effects and corrected for individual differences across subjects by computing the RTPMs. **Panels E** and **F** of **Figure 8** show the mean RTPM results for active and sham conditions, respectively.

**Figure 8.**
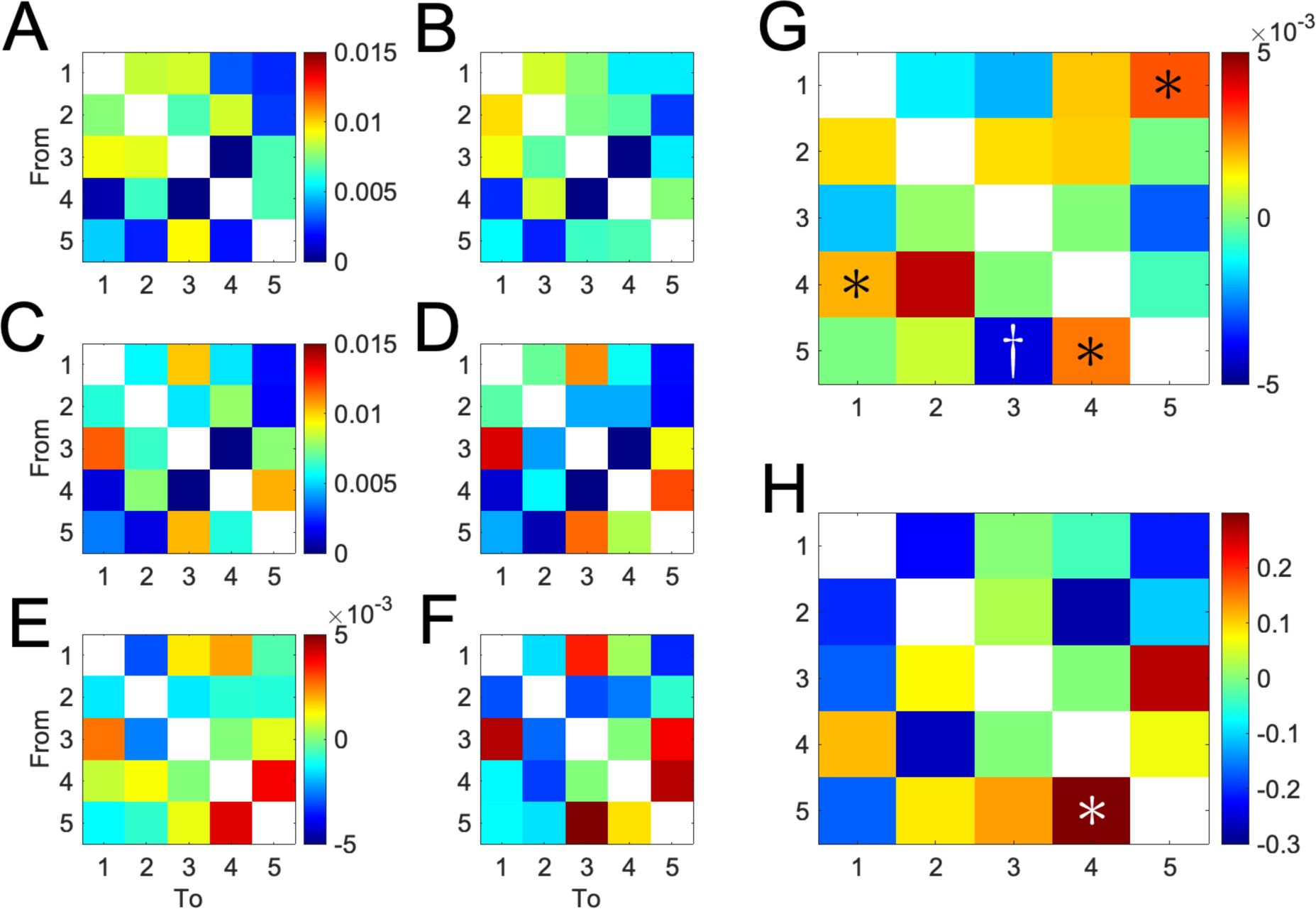
Transition probability matrices (TPMs), relative transition probability matrices (RTPMs), and correlations with LC MTC. Each colored square in **(A)-(F)** shows the transition probability from state *i* to state *j*, with *i* represented by rows and *j* represented by columns. **(A)** and **(B)** display the TPMs for RS0 in the active and sham conditions, respectively. **(C)** and **(D)** show the TPMs for PostAr for active and sham conditions, respectively. **(E)** and **(F)** show the RTPMs for active and sham conditions, i.e. the differences between TPMs for PostAr and RS0 (**(E)** = **(C)** – **(A)** and **(F)** = **(D)** – **(B)**). **(G)** shows that several of the differences in transition probability from baseline were significantly larger in the active than the sham condition (S4→S1, S5→S4, and S1→S5; **(G)** = **(E)** – **(F)**, Wilcoxon sign rank test against 0), while one difference in transition probably from baseline was trending smaller (S5→S3). Finally, **(H)** shows the Spearman correlation coefficients relating values shown in **(F)** and LC MTC across subjects; one relative transition probability (S5→S4) reached significance. See main text for details. * p < 0.05; ^†^ p < 0.1. Here we do not correct for multiple comparisons, following practices in MRI research to control family-wise error rate under planned statistical testing.

To address how these transition probabilities relative to baseline changed as a function of active squeeze, we examined the Euclidean distance between these RTPMs for each subject (RTPM_Active_ – RTPM_Sham_, **Figure 8G**; see **Methods Section 2.3.2.2**). This process identified that on the whole, the distances between RTPM_Active_ and RTPM_Sham_ were significantly different from 0 across subjects (Wilcoxon sign rank, Mann-Whitney U = 465, p = 1.7344e-06), indicating that the change in transition probabilities from RS0 to PostAr was different overall in active versus sham conditions. In other words, the transition probabilities revealed differences between active and sham that were obscured by the average state duration analysis described above.

##### 3.3.2.2 State-specific analysis of transition probabilities

However, the significant difference in overall RTPMs between active and sham does not account for the transitions between specific states. To test whether state-specific changes in transition probabilities from baseline were different between active and sham conditions, we conducted separate Wilcoxon sign rank tests for each state in the off-diagonal transition probabilities in RTPM_Active_ – RTPM_Sham_; that is, for all off-diagonal transition probabilities going from state *j* to state *k*, we computed the difference, RTPM_Active,i,j,k_ – RTPM_Sham,i,j,k_ (*i*∊{*subjects*}) and then tested whether the distribution of these differences across subjects was significantly different from 0 for every pairwise state transition using a Wilcoxon sign rank test. Now, we can see which pairwise state transitions are likely to be driving the overall differences between active and sham RTPMs. Several states showed significant differences between RTPM_Active_ and RTPM_Sham_ that were specific to certain state transition probabilities, summarized in **Figure 8F**.

Here, we focus on transitions into and out of S4, as that is the SN-dominant state which we found above to have duration significantly correlated with LC MTC regardless of active versus sham condition. In the RTPM differences (RTPM_Active_ – RTPM_Sham_; **Figure 8G**), we now see that two transitions that involve S4 were significantly different between active and sham conditions after correcting for baseline effects. Namely, S4→S1 (p = 0.0467) and S5→S4 (p = 0.0226) were both significantly more likely to occur in the active than the sham condition.

##### 3.3.2.3 Relationships to LC integrity

Finally, we were interested in how LC MTC were related to changes in transition probabilities from RS0 to PostAr, especially given the observation that LC MTC significantly correlated with S4 duration in the previous analysis. To answer this question, LC MTC values were correlated with RTPM_Active_ – RTPM_Sham_ across all subjects prior to obtaining the global average (**Figure 8G**). Once again, we saw significant correlations having to do with S4, the SN-dominant state. A significant correlation was observed between LC MTC and the changes in the S5→S4 transition from baseline (p = 0.0150), indicating that LC MTC levels may relate to the probability of subjects transitioning out of the whole-brain deactivation state and into the SN-dominant state as a function of the handgrip task relative to baseline.

Together, these results paint a compelling picture. LC MTC appears to be related to the persistence of an SN-dominant state (S4) once entered, regardless of how the subject ended up in the state, as well as the propensity to transition into that state, especially from positions of relative overall deactivation. However, one potential confounding factor is that LC MTC might simply correlate with the predominance of S4 in general: If S4 is more likely to occur in subjects with higher neuronal density in LC regardless of task or previous state, that could offer a less exciting explanation for why we have seen the patterns described here. We sought to rule out this possibility by conducting the fractional occupancy analyses described next.

#### 3.3.3 Fractional occupancy

To determine whether the results above may be trivially due to differences in fractional occupancy in S4 (or other states) across active and sham conditions, or differences in LC MTC that drove such a propensity to occupy S4, we directly examined the percent of time each subject spent in each state. That is, we calculated the fractional occupancy (see **Methods Section 2.3.2.3**) for each state and within each block (RS0 vs PostAr), and then performed analyses similar to those done above for average state duration on the difference between these, Δ*FO*.

The 2 (condition: active vs sham) x 5 (state) repeated measures ANOVA revealed a main effect of state (F(4,100) = 14.51, p < .001) and a trending main effect of condition (F(1,100) = 3.90, p = 0.059), but no interaction (F(4,100) = 1.73, p = 0.150) (**Figure 9A**). We thus collapsed across conditions and conducted a one-way repeated measures ANOVA to evaluate whether state affected fractional occupancy regardless of active versus sham condition. This approach revealed a main effect of state (F(4,116) = 14.27, p < .001), so we followed these findings with five t-tests against 0 to discover which states deviated in their fractional occupancy between RS0 and PostAr. These tests revealed that subjects spent a significantly higher percentage of their time in S2 during the PostAr block versus RS0, but significantly lower percentage of the time in S3 and S4 (**Table 7**). While these may seem surprising, recall that fractional occupancy is defined as the percentage of time spent in a particular state relative to the total length of the scan, and the RS0 block is only 150 TRs long while the PostAr block is 375. Thus, any occupancy in short-lived states like S4 (as discussed above) during RS0 might lead to a relatively outsized appearance of fractional occupancy.

**Figure 9.**
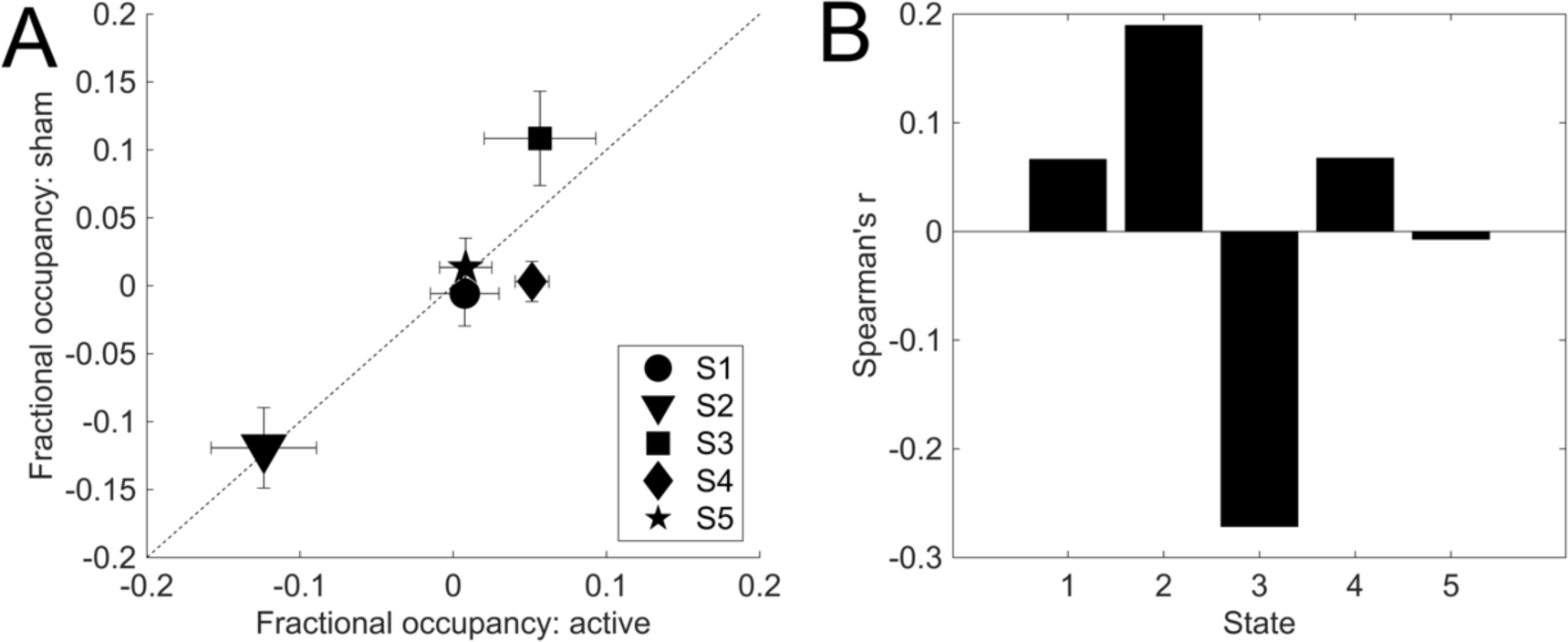
Average baseline-corrected fractional occupancy (Δ*FO*) in the active and sham conditions, and correlations with LC MTC across subjects. **(A)** The 2 (condition: active vs sham) x 5 (state) repeated measures ANOVA revealed no main effect of condition but a main effect of state, which is visible in the nearness of the points to the identity line. **(B)** No state’s fractional occupancy significantly correlated with LC MTC. Error bars in **(A)** show the standard error of the mean. See main text for details.

**Table 7.** Average fractional occupancy and t-test results for comparing fractional occupancy changes from baseline across state, collapsed across condition. Three states (S2, S3, and S4) showed significant differences in fractional occupancy during PostAr relative to baseline (RS0): S2 was occupied significantly more often in PostAr, while S3 and S4 were occupied significantly less often relative to the length of the block. See main text for details. * p < 0.05. All significant effects survive correction for multiple comparisons via the Benjamini-Hochberg method.

| | Mean $\Delta FO$ (PostAr-RS0) | $\sigma$ | t | p |
| --- | --- | --- | --- | --- |
| **State 1** | -0.0009 | 0.0937 | -0.0506 | 0.960 |
| **State 2** | 0.1215 | 0.1103 | 6.0354 | < .001* |
| **State 3** | -0.0825 | 0.1262 | -3.5827 | 0.001* |
| **State 4** | -0.0273 | 0.0511 | -2.9240 | 0.007* |
| **State 5** | -0.0108 | 0.0878 | -0.6754 | 0.505 |

Nevertheless, our goal with this analysis was to identify a potential origin for the aforementioned result demonstrating that average duration of S4, or transition into S4 or other states, was related to LC MTC. Thus, as a final check we correlated Δ*FO* with LC MTC across subjects. This analysis revealed no significant correlations between the fractional occupancy in any state and LC MTC (**Figure 9B**; **Table 8**). We can therefore be confident that the relative higher probability of transitioning S5→S4 in the active condition, and the observed correlation between this specific transition and LC MTC across subjects, is unlikely to be attributed to percent of time spent in S4 or any effect of LC MTC on this fractional occupancy. Instead, it seems likely that the S5→S4 transition as well as the duration of S4 – and their correlations with LC MTC – may reflect a core role of LC neuronal density on the effectiveness of the active squeeze manipulation on the ease of entering into an aroused state.

**Table 8.**
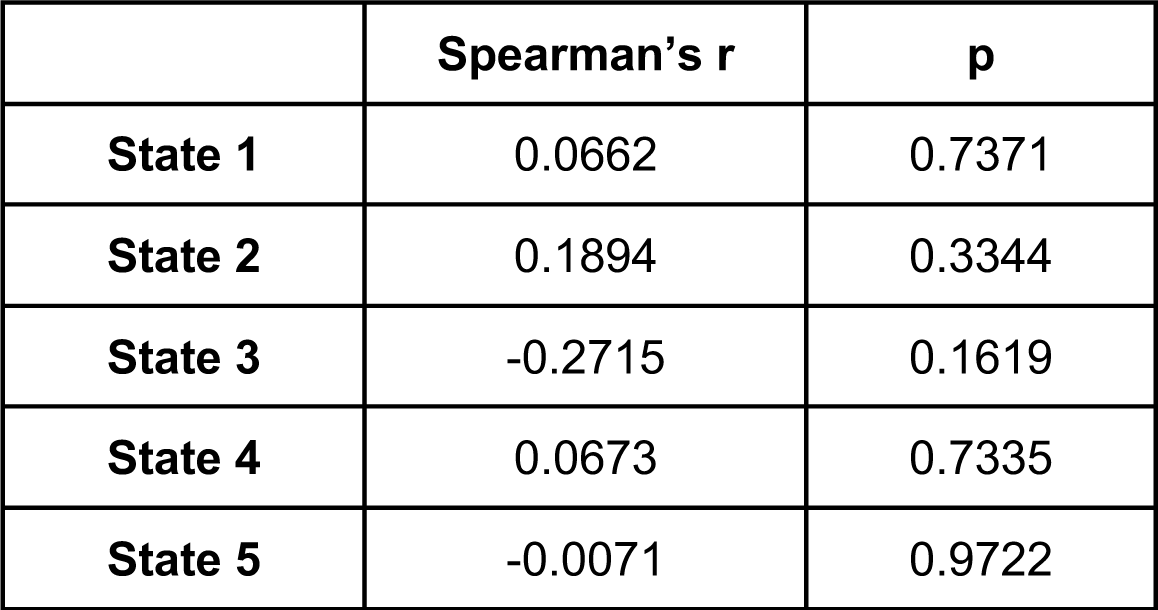
. Spearman correlation coefficients comparing LC MTC to Δ*FO* (average fractional occupancy change from baseline), collapsed across conditions. No changes in fractional occupancy were significantly correlated with LC MTC. See main text for details.

|  | **Spearman's r** | **p** |
| --- | --- | --- |
| **State 1** | 0.0662 | 0.7371 |
| **State 2** | 0.1894 | 0.3344 |
| **State 3** | -0.2715 | 0.1619 |
| **State 4** | 0.0673 | 0.7335 |
| **State 5** | -0.0071 | 0.9722 |

Finally, we examined the correlation between average state duration and fractional occupancy across subjects as a function of state. We computed the Pearson correlation between these metrics for each state (**Table 9**).

**Table 9.** Correlation between fractional occupancy and average state duration across subjects as a function of state. * p < 0.05. All significant effects survive correction for multiple comparisons via the Benjamini-Hochberg method.

|  | **Pearson's r** | **p** |
| --- | --- | --- |
| **State 1** | 0.7345 | < .001* |
| **State 2** | 0.6654 | < .001* |
| **State 3** | 0.6099 | < .001* |
| **State 4** | 0.4705 | 0.012* |
| **State 5** | 0.4894 | 0.008* |

### 3.4 Discussion

#### 3.4.1 Summary and interpretation of results

Here, we sought to characterize the consequences of LC up-regulation on the spatiotemporal dynamics of brain networks. We fitted a hidden Markov model (HMM) to whole-brain fMRI data while healthy adult humans performed a pseudo-resting-state task designed to engage LC, and examined how LC activation modulated brain states themselves as well as the dynamics of transitions between them. The HMM identified five stable states corresponding to patterns of activity in the default mode network (DMN), dorsal attention network (DAN), front-parietal control network (FPCN), salience network (SN), and the LC. One of these states, the SN-dominant state – which occurred during active engagement with the squeezing task (during both active and sham conditions) – was highly stable across conditions, although other states showed significant difference between active and sham. Interestingly, the stability of this SN-dominant state, S4, was correlated with LC MTC.

We also found that S4 showed intriguing and asymmetrical transition probabilities as a function of active versus sham squeeze. Specifically, when subjects actively squeezed, they were more likely to go from a relatively deactivated state (S5) to an SN-dominant state (S4) than when they simply brought their hand to their chest, and likewise that once they achieved that SN-dominant state (S4) they were more likely to transition into a DMN-dominant state (S1) in the active than in the sham condition (**Results Section 3.3.2.2**). This suggests not only that the squeeze manipulation was successful in easing the transition into an SN-dominant state (Hussain et al., 2019; Kozłowski et al., 1973; Lake et al., 1976; Mather et al., 2020; Nielsen et al., 2015; Nielsen & Mather, 2015; Wallin et al., 1992, 1987), but also that once the SN-dominant state was achieved, subjects more easily transitioned into an internally-oriented mental state (DMN is associated with internally-oriented attention and self-referential mind wandering (Raichle, 2011, 2015)) than in the sham condition.

This pattern might seem counterintuitive, as one might expect an attentional state (e.g. S2) being the more likely transition after arousal. One possible explanation is that NE depletion played a role, in that the arousing stressor (squeezing) depleted the LC’s supply of NE causing subjects to return to the DMN-dominant state rather than switching into an attention-dominant state (i.e., S2), thereby prompting an attention “reset” (Mather et al., 2016; Sara, 2015, 2016) rather than a direct transition into S2. Moreover, recall that, aside from the squeeze/hand-raising task, there was no other task for the subjects to do in the scanner; therefore, after entering the SN-dominant state in the active condition, when it started to dissipate they may have had nothing external to focus on and therefore focused on their own thoughts more than in the sham condition. This interpretation is consistent with the observation that the S1→S5 transition was also significantly more likely to occur in active than in sham after correcting for baseline effects (p = 0.0168). Interestingly, this suggests the presence of a circular S1→S5→S4→S1 loop that is more likely to occur in active than in sham, but which is broken by the (trending; p = 0.0937) stronger presence of S5→S3 in the sham condition (recall that S3 is a more general “everything is active” state rather than the internally-focused S1 or the externally-focused S2). Recall that negative values in **Figure 8G** indicate instances where the change in transition probability relative to baseline was lower during the active condition than in the sham condition. Thus, the trending negative value of S5→S3 occurred because the active squeeze impeded an increase in this transition probability relative to baseline, rather than the active squeeze having no or a reduction effect on this probability. This suggests a difference in LC activity between the active squeeze versus the sham (bring arm to chest) control: While lifting the arm to the chest may marginally up-regulate LC activity and induce some arousal (Hussain et al., 2019), there is also a relatively high probability of transitioning between whole-brain activation and whole-brain deactivation states in pure resting state (i.e., no actual task) paradigms (S. Chen et al., 2016). These results are consistent with previous findings showing common back and forth transitions between whole-brain activation states and whole-brain deactivation states in the absence of a task (S. Chen et al., 2016). It is possible that the transition S5→S3 was trending significant but S3→S5 was not because slight changes in LC activity led to an increased probability of transitioning out of a whole-brain deactivation state (S5) and into a whole-brain activation state (S3) where attentional networks were activated, rather than the other way around (Hussain et al., 2019).

Although we can only speculate here as to why this S1→S5→S4→S1 loop was more likely to occur in active than in sham, it appears that the increased arousal due to the active squeeze may have kept participants more engaged throughout the scan, even if that engagement was oriented inwards. It is of course also possible that subjects transitioning out of the DMN-dominant state and into the whole-brain deactivation state could indicate that a network not analyzed in this study is at work in response to the active squeeze; we leave the exploration of this possibility to future studies.

Interestingly, no subject experienced S3→S4 or S4→S3 transitions in either the active or sham condition (darkest blue squares in **Figure 8A-D**). To understand the significance of this observation, recall that S3 is the “everything is activated” state and S4 is the SN-dominant state. One possible explanation for this observation is that LC may be involved in allocating attention (Sara, 2015). Previous studies have shown that an increase in LC activity may cause a rapid shift in the allocation of attention in response to the sudden onset of a stimulus (Sara, 2015, 2016). This may be why subjects did not transition from the SN-dominant state (S4) into the whole-brain activation state (S3) where all attention-related networks were concurrently activated, and the LC induced some phenomenon that could not be observed in this investigation. Instead, transitions from S4 into S1, S2, or S5 were seen where DMN, FPCN, DAN, and SN were not synchronously activated – only a few (two at most as seen in S2) were activated at the same time. The LC allocated attention to DAN and SN since the subjects were observed to transition out of S4 into S2. Attention was also allocated to a resting state network (DMN in S1) since some subjects returned to resting state following the squeeze periods. Some attention could have been allocated to a network not examined (S5), albeit more so in the sham condition than in the active, but never to DMN, FPCN, DAN, and SN at the same time. Because this idea of attentional allocation is associated with network resetting, our results may provide preliminary evidence that HMM-derived states can test the LC network reset hypothesis (Sara, 2015, 2016). However, additional tests focusing on attentional shift during these specific transitions are needed.

Finally, we also found that the degree to which the squeeze task affected subjects’ propensity to transit into S4 was also correlated with LC MTC, and LC MTC also showed correlations with the duration of that state once it was achieved, regardless of active versus sham condition. In aggregate, we interpret these results to suggest that, even in healthy young adults, LC structure may play an important role in facilitating the transition into a state characterized by SN engagement, moderating the efficacy of any task or condition that might engage the LC circuit to cause norepinephrine release.

#### 3.4.2 Future directions and limitations

These findings can be applied in future studies to better characterize previous observations that LC activation contributes to not just an aroused state but also a U-shaped function relating arousal and performance, known as the Yerkes-Dodson curve (Aston-Jones & Cohen, 2005). Our data-driven approach, using a HMM to reveal hidden states of network activity patterns, clearly identified an SN-dominant state in S4 which was highly stable even in the absence of an active stress-type manipulation (the squeeze), but which nevertheless showed meaningful hallmarks both of our behavioral manipulation and of individual differences in LC integrity. In particular, under the active squeeze condition, network nodes in SN exhibited stronger activation and subjects tended to persist longer in this state when they actively squeezed, depending on their LC MTC. Notably, these results demonstrate that network-based approaches to studying the effects of LC up-regulation can identify meaningful patterns even in the absence of precise measurement of LC activity itself. That the squeeze manipulation also eased the transition into this SN-dominant state from a state of relative deactivation is not only consistent with previous findings showing that attention-related states specifically related to stimulus salience may be more easily achieved if LC is successfully engaged (Hussain et al., 2019; Kozłowski et al., 1973; Lake et al., 1976; Mather et al., 2020; Nielsen et al., 2015; Nielsen & Mather, 2015; Wallin et al., 1992, 1987), but also allow us to examine what that state of salience-sensitivity consists of: relative deactivation of nearly every other network in the brain studied here with the exception of two nodes in FPCN (left and right posterior inferior parietal lobule) and two nodes in DAN (left anterior prefrontal cortex and left anterior insula). In contrast, DMN was almost entirely deactivated in this SN-dominant state, in stark contrast to its high degree of activation that likely occurred directly after presence in the SN-dominant state (in the DMN-dominant state, S1). Moreover, that the S4→S1 transition was more likely in the active than sham condition – and that the subsequent S1→S5 transition was also more likely – suggests a cycle of outward orientation (S4) followed by relatively inward-directed attention (S1) and then a general resetting (S5) that is significantly associated with LC activity levels. Future research may more explicitly explore these cycles and include more brain networks to more fully characterize recurring patterns of transitions through state space and their relationship to LC activity and integrity, as well as using this approach to study the extreme ends of the U-shaped Yerkes-Dodson curve (Aston-Jones & Cohen, 2005).

It is also worth discussing the relationship between the results found here and a more general definition of ‘arousal’, broadly construed, in the context of effort. Specifically, there is a growing body of evidence relating LC production of norepinephrine to effort and mobilization of resources. For example, firing of noradrenergic neurons has been related to effort production in relation to the energization of behavior to face the challenge at hand (Varazzani et al., 2015). Likewise, it has been hypothesized that the multiple effects of noradrenaline manipulation on behavior could be captured by a specific modulation of a single ‘effort’ variable identifiable through computational modeling (Borderies et al., 2020), also providing a potential avenue for intervention in this pathway in modifying pathological perceptions of effort for clinical populations with LC degeneration (Hezemans et al., 2022). Given that our task intervention in this study consisted of a hand-grip task that required effort production in order to perform, it is also possible that our results might speak to brain networks involved in perceived and actual effort as a function of LC MTC in healthy young adults as well. As we did not collect perceived effort measures here, however, we must leave exploration of these interesting questions to future studies that may follow up on the methods and results presented here.

Although robust across several metrics, we do note that the results described here are a first step in a longer research program, given that the effects we found here were relatively small in size. This may have occurred because the handgrip manipulation was not strong enough to induce changes in LC activation (and consequences in brain states and dynamics) large enough to be detected via fMRI. Future studies may wish to employ a more aggressive manipulation, such as subjects dipping their hand in cold water, administering electrical pulses, or presenting jarring sounds (Marmon & Enoka, 2010; Oyarzún et al., 2012; Redondo et al., 2008; Schwabe & Schächinger, 2018; Stark et al., 2006). It is also possible that we would see stronger correlations between functional connectivity and contrast in older adults or in a population with disease or neurodegeneration (e.g., Alzheimer’s, Parkinsons) as contrast has been shown to vary in both populations (Hwang et al., 2023; Liu et al., 2019; Ohtsuka et al., 2013; Sasaki et al., 2006; Shibata et al., 2006). Nevertheless, we did observe reasonable variation in LC MTC across subjects even in this young, healthy population, contributing to knowledge of individual differences in LC structure and function. Future studies using these other manipulations or more precise measurements of LC activity – perhaps in nonhuman animals – may be able to shed more light on these questions, and the effect sizes reported here can serve to facilitate power analyses for such future research as well.

On the other hand, if the squeeze manipulation we used did in fact induce a robust LC response and consequences in brain dynamics, it is also possible that the measures used in this investigation were not sensitive enough to detect it due to poor signal-to-noise ratio in the BOLD signal. Future investigations could explore various denoising methods for BOLD data (Kundu et al., 2012; Quian Quiroga & Garcia, 2003). As mentioned above, LC BOLD is subject to physiological noise, specifically cardiac pulsation and respiration (Clewett et al., 2016; Glover et al., 2000; K. Y. Liu et al., 2017; Mather et al., 2017). Here, we were unfortunately unable to regress out these potential noise sources because physiological data were not collected in conjunction with functional data. Future studies should plan to collect cardiac pulsation and respiration data so that physiological noise can be regressed out from the LC BOLD signal prior to HMM fitting (Glover et al., 2000).

Finally, it is worth noting that we did not explore other HMM variants, including previous approaches that have explored fitting HMMs to state-specific covariance matrices (D. Vidaurre et al., 2021; Diego Vidaurre, 2021; Diego Vidaurre, Hunt, et al., 2018), or using sliding window approaches to compute functional connectivity and then fitting HMMs to the resulting correlation matrices among nodes in various brain networks (Hussain et al., 2022; Ou et al., 2015). Here we elected to focus on activity-based HMMs, as our main question of interest was in demonstrating a first step towards characterizing activity-dependent state changes as a function of structural changes in LC and/or its activity levels. While a full exploration of the many covariance-based HMM approaches is beyond the scope of this manuscript, we encourage future work to follow up on the insights that functional-connectivity or covariance HMMs might add to the LC-dependent state changes identified in the present results.

#### 3.4.3 Conclusions

Overall, we have shown that individual differences in LC MTC – even in healthy young adults – are associated with the stability of an arousal-modulated brain state dominated by salience network activity, and the propensity to enter and dwell in this state. Our results reveal important new insight into the role of LC in relation to the dynamics of brain states related to arousal, stimulus sensitivity, and task engagement.

## Data/code availability statement

Hidden Markov models were generated using the hmmlearn library in python (https://github.com/hmmlearn/hmmlearn). The fMRI dataset and pupillometry data are available on Dryad at https://doi.org/10.7280/D1HQ3B.

## Authorship contribution

**<u>Sana Hussain:</u>** Conceptualization, Formal Analysis, Investigation, Methodology, Project Administration, Software, Validation, Visualization, Writing –– Original Draft, Writing –– Review & Editing. **<u>Isaac Menchaca:</u>** Investigation, Methodology, Formal Analysis. **<u>Mahsa Alizadeh</u> <u>Shalchy:</u>** Investigation, Methodology. **<u>Kimia Yaghoubi:</u>** Investigation, Methodology. **<u>Jason</u> <u>Langley:</u>** Investigation, Methodology, Supervision, Validation, Visualization, Writing –– Review & Editing. **<u>Aaron R. Seitz:</u>** Conceptualization, Funding Acquisition, Investigation, Methodology, Project Administration, Supervision, Validation, Visualization, Writing –– Review & Editing. **<u>Xiaoping P. Hu:</u>** Conceptualization, Funding Acquisition, Investigation, Methodology, Project Administration, Resources, Supervision, Validation, Visualization, Writing –– Review & Editing. **<u>Megan A. K. Peters:</u>** Conceptualization, Funding Acquisition, Investigation, Methodology, Project Administration, Resources, Supervision, Validation, Visualization, Writing –– Original Draft, Writing –– Review & Editing.

## Acknowledgements

This work was supported in part by the UCR NASA MIRO FIELDS Fellowship (to Sana Hussain). Data collection and sharing for this project was provided by the Human Connectome Project (HCP; Principal Investigators: Bruce Rosen, M.D., Ph.D., Arthur W. Toga, Ph.D., Van J. Weeden, MD). HCP funding was provided by the National Institute of Dental and Craniofacial Research (NIDCR), the National Institute of Mental Health (NIMH), and the National Institute of Neurological Disorders and Stroke (NINDS). HCP data are disseminated by the Laboratory of Neuro Imaging at the University of Southern California. This work was funded by NIA R01 NS108638-01 (PIs: Xiaoping P. Hu and Aaron R. Seitz) and by the Canadian Institute for Advanced Research Azrieli Global Scholars Program (PI: Megan A. K. Peters). Funding sources had no involvement in the design and methodology of the study.

## Supplementary Material

### S1. Regions of interest

Table S1 shows the labels, and MNI coordinates for all networks and ROIs discussed and were used to center a 5mm^3^ isotopic marker, except for the LC ROIs (which was derived from an atlas and whose voxels were split into rostral and caudal regions) (Deshpande et al., 2009, 2011; Stilla et al., 2007). The BOLD signal from each voxel within an ROI were extracted and averaged to represent the overall signal for an ROI. This was repeated for 31 total ROIs: 9 from DMN, 7 from FPCN, 6 from DAN, 7 from SN, and 2 from LC. Although the LC is only a single ROI split into rostral and caudal portions, it is an entity distinguishable from the large-scale networks and will henceforth be referred to as a network in this paper.

**Table S1.** List of MNI coordinates used for ROIs in the default mode network (DMN), fronto-parietal control network (FPCN), dorsal attention network (DAN), salience network (SN), and locus coeruleus (LC). Talaraich coordinates for DMN, FPCN, and DAN were taken from Deshpande et al. (Deshpande et al., 2011) and were converted to MNI using (Brett et al., 2002; Deshpande et al., 2011; Laird et al., 2005; Lancaster et al., 2007) while MNI coordinates for SN were taken directly from Raichle 2011 (Raichle, 2011). MNI coordinates for LC were taken from (Langley et al., 2020).

| ROI | Abbrev. | Full Name | MNI (x,y,z) | Source |
| --- | --- | --- | --- | --- |
| **Default Mode Network** |  |  |  |  |
| 1 | PCC | Posterior Cingulate Cortex | (2,54,16) | Deshpande et al. 2011 |
| 2 | L pIPL | Left Posterior Inferior Parietal Lobule | (-46,-72,28) | Deshpande et al. 2011 |
| 3 | R pIPL | Right Posterior Inferior Parietal Lobule | (50,-64,26) | Deshpande et al. 2011 |
| 4 | OFC/vACC | Orbitofrontal Cortex/Ventral Anterior Cingulate Cortex | (4,30,26) | Deshpande et al. 2011 |
| 5 | dMPFC BA 8 | Dorsomedial Prefrontal Cortex Brodmann Area 8 | (-14,54,34) | Deshpande et al. 2011 |
| 6 | dMPFC BA 9 | Dorsomedial Prefrontal Cortex Brodmann Area 9 | (22,58,26) | Deshpande et al. 2011 |
| 7 | L DLPFC | Left Dorsolateral Prefrontal Cortex | (-50,20,34) | Deshpande et al. 2011 |
| 8 | L PHG | Left Parahippocampal Gyrus | (-10,-38,-2) | Deshpande et al. 2011 |
| 9 | L ITC | Left Inferolateral Temporal Cortex | (-60,-20,-18) | Deshpande et al. 2011 |
| **Fronto-Parietal Control Network** |  |  |  |  |
| 1 | L aPFC | Left Anterior Prefrontal Cortex | (-36,56,10,) | Deshpande et al. 2011 |
| 2 | R aPFC | Right Anterior Prefrontal Cortex | (34,52,10) | Deshpande et al. 2011 |
| 3 | R DLPFC | Right Dorsolateral Prefrontal Cortex | (46,14,42) | Deshpande et al. 2011 |
| 4 | L aINS | Left Anterior Insula | (-30,20,-2) | Deshpande et al. 2011 |
| 5 | R aINS | Right Anterior Insula | (32,22,-2) | Deshpande et al. 2011 |
| 6 | L aIPL | Left Anterior Inferior Parietal Lobule | (-52,-50,46) | Deshpande et al. 2011 |
| 7 | R aIPL | Right Anterior Inferior Parietal Lobule | (52,-46,46) | Deshpande et al. 2011 |
| **Dorsal Attention Network** |  |  |  |  |
| 1 | L MT | Left MidThalamus | (-44,-64,-2) | Deshpande et al. 2011 |
| 2 | R MT | Right MidThalamus | (50,-70,-4) | Deshpande et al. 2011 |
| 3 | L FEF | Left Frontal Eye Field | (-24,-8,50) | Deshpande et al. 2011 |
| 4 | R FEF | Right Frontal Eye Field | (28,-10,50) | Deshpande et al. 2011 |
| 5 | L SPL | Left Superior Parietal Lobule | (-26,-52,56) | Deshpande et al. 2011 |
| 6 | R SPL | Right Superior Parietal Lobule | (24,-56,-54) | Deshpande et al. 2011 |
| **Saliience Network** |  |  |  |  |
| 1 | DAC | Dorsal Anterior Cingulate | (0,-22,36) | Raichle 2011 |
| 2 | L aPFC | Left Anterior PFC | (-34,44,30) | Raichle 2011 |
| 3 | R aPFC | Right Anterior PFC | (32,44,30) | Raichle 2011 |
| 4 | L Insula | Left Insula | (-40,2,6) | Raichle 2011 |
| 5 | R Insula | Right Insula | (42,2,6) | Raichle 2011 |
| 6 | L LP | Left Lateral Parietal | (-62,-46,30) | Raichle 2011 |
| 7 | R LP | Right Lateral Parietal | (62,-46,30) | Raichle 2011 |
| **Locus Coeruleus** |  |  |  |  |
| 1 | R LC | Rostral locus coeruleus | left (-3.0, -36.7, -20.4)<br>right (4.4, -36.4, -19.5)<br>(Probabilistic atlas) | Langley et al. 2020 |
| 2 | C LC | Caudal locus coeruleus | left (-5.9, -38.1, -29.9)<br>right (7.0, -38.1, -30.3)<br>(Probabilistic atlas) | Langley et al. 2020 |

### S2. Supplemental methods and results

#### S2.1 Determining model order

We examined the stability of model orders 3-15 (range based on previously established norms; (S. Chen et al., 2016; Hussain et al., 2022; Z. Yang et al., 2008)) through two methods. First, we adopted the Ranking and Averaging Independent Component Analysis by Reproducibility (RAICAR) method (S. Chen et al., 2016; Hussain et al., 2022; Z. Yang et al., 2008) which computes Pearson correlations among states recovered across three initializations of the HMM fitting procedure: one with uniform starting probability of residing in all states, and two with randomly assigned starting probabilities. Note that, because within a given initialization the labeling of each state as “state 1” or “state 2” is arbitrary, in order to assess whether the recovered states are actually the same across initializations, we must first ‘match’ them up. We accomplished this matching via Pearson correlations, such that e.g. state 1 from Initialization 2 was relabeled as state 2 just in case the Pearson correlation between that state and state 2 from Initialization 1 was higher than any other pairwise correlation. Thus, after relabeling, each state label across initializations universally corresponded to the same spatial pattern to the maximal extent possible. Within each state assignment, the matched state patterns were then Pearson correlated between all initializations to obtain (# *states*)!/2! * (#*states* − 2)! Pearson’s R values, thereby determining the maximal degree of similarity between the matched patterns for the model order being currently tested. Finally, these values were then averaged, sorted from largest to smallest, and plotted as a function of model order, and then compared to a critical threshold of R = 0.9. That is, for a given state to be considered ‘stable’ across those initializations of the model, we must be able to discover a pairing of (arbitrarily-labeled) states across those two initializations such that the correlation between the two states is 0.9 or higher. Previously, a stability threshold of 0.8 has been used when more ROIs are being tested (S. Chen et al., 2016; Z. Yang et al., 2008), but here we examined only 31 ROIs and so opted for a more conservative threshold of 0.9. Model orders for which several states displayed ‘stability’ less than R = 0.9 were considered unstable, i.e. too few or too many hidden states specified *a priori*. See the methods described by Hussain and colleagues (Hussain et al., 2022) for more detail.

As a second, confirmatory approach, we also employed Euclidean distances to assess model stability and model order. In this case, a smaller Euclidean distance between two matched states indicates a better correspondence between model initializations. We followed similar logic to that used in the RAICAR method, but with additional steps to ensure that states from Initialization 2 were optimally matched with states from Initialization 1 and then relabeled, and the same for Initialization 3. To do this, state assignments from two initializations, I_i_ and I_j_, within a certain model order were permuted and their spatial patterns matched via the smallest Euclidean distance such that each state universally corresponded to the same spatial pattern. For example, after permuting the state assignments from I_i_ and I_j_, the Euclidean distance between state 1 from I_i_ and all states from I_j_ are computed. The results may show that the smallest Euclidean distance was computed with state 5 from I_j_ indicating that state 1 from R_i_ best corresponds to state 5 from R_j_. State 5 from I_j_ would thus be relabeled as state 1 from I_j_, then the process is repeated between state 2 from R_i_ and all states from I_j_ (except the relabeled state 1 as it has already been matched). This permutation-and-matching procedure was performed 100 times for all pairs of realizations (where matching I_i_ → I_j_ is not distinguished from matching I_j_ → I_i_) to ensure that the spatial patterns are paired uniquely without any bias of state assignment. This was repeated for a range of model orders generating a total of 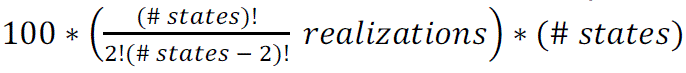 values for a particular model order which are then averaged to represent its overall stability. This single average was plotted as a function of model order producing a curve where the smallest value succeeded by continuously increasing values for higher model orders indicates the optimal number of states for a dataset. Although this may seem similar to the RAICAR-based method, the Euclidean distance based method is more conservative because stability is assessed by ensuring that the average of hundreds of Euclidean distances is below a prespecified threshold rather than the average of a handful of R values.

#### S2.2 Confirmation that active squeeze engages LC

We checked to confirm that the active squeeze condition was associated with increased LC BOLD signal than was the sham control condition by computing LC BOLD signal in rostral and caudal regions of the LC during the RS3 period in comparison to the RS0 baseline for each subject. That is, for each subject and within each LC region (rostral and caudal) we subtracted average BOLD signal during RS0 from BOLD signal during RS3 at each TR, and plotted the timecourse of these fluctuations averaged across subjects. This analysis revealed that the active squeeze condition produced a sustained increase in LC BOLD relative to RS0 baseline for the duration of the RS3 period, demonstrating that active squeeze was successful in increasing LC activity (**Figure S1**).

**Figure S1.**
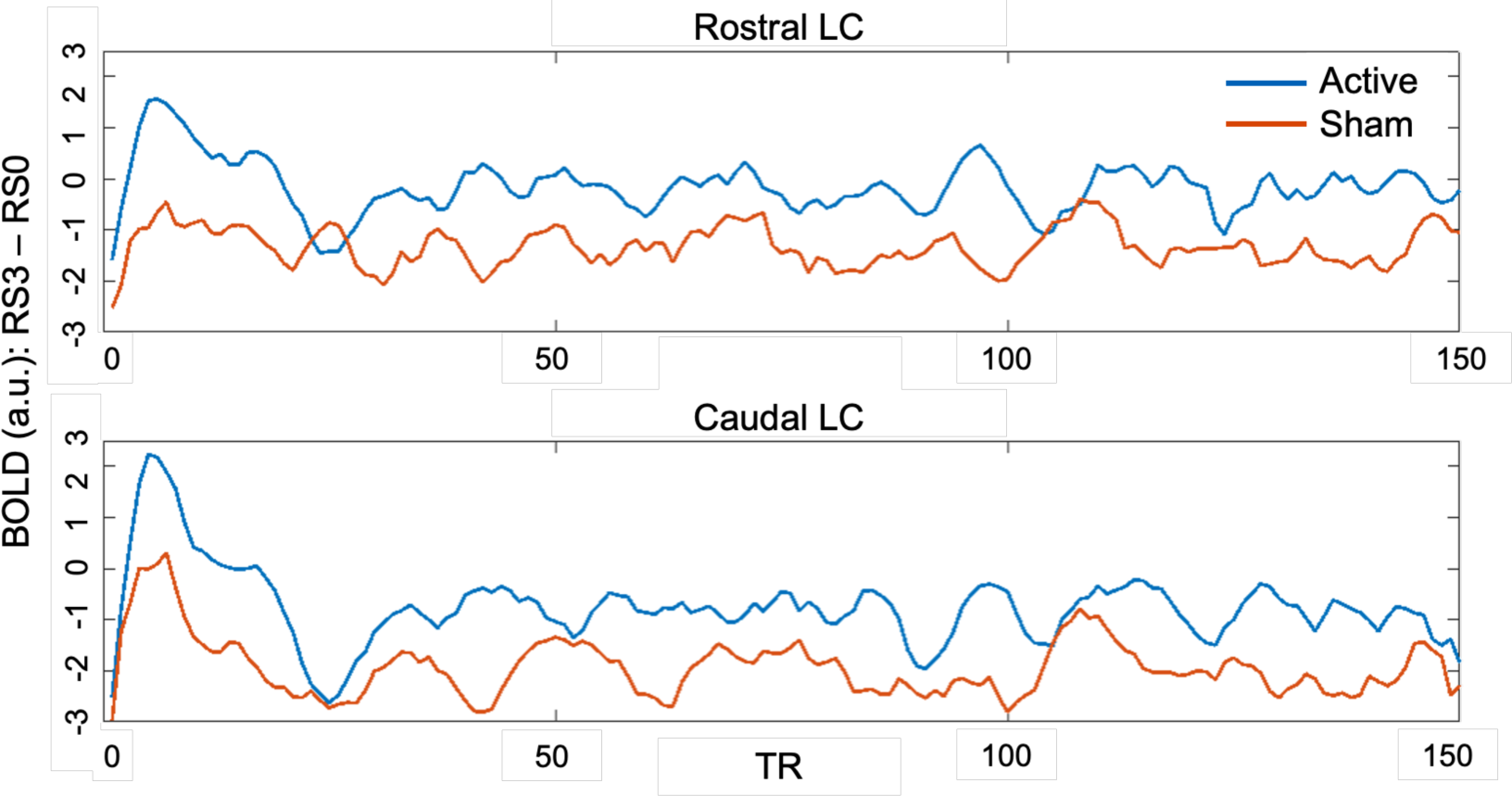
Active squeeze versus sham control effects on LC BOLD signal during the RS3 period, relative to baseline (RS0). Both rostral and caudal LC regions demonstrated a sustained increase in BOLD signal compared to baseline during this period for the active squeeze over sham control, confirming that the squeeze manipulation successfully increased LC activity.

### S3. Pupillometry

#### S3.1 Pupillometry methods

We also collected simultaneous pupillary dilation as a proxy for LC activity. Fluctuations in pupil diameter have been validated in animal models using invasive recordings (Joshi et al., 2016), allowing them to be used as a noninvasive proxy measure for LC activity when simultaneously recorded with fMRI in humans (Joshi et al., 2016; Murphy et al., 2014). As noted below, however, technical challenges during data collection and relatively sparsity of data due to stringent exclusion requirements precluded strong conclusions being drawn via the pupillometry data and analyses. For completeness, here we present the exploratory analyses we were able to conduct, and show how they qualitatively align with the main findings.

##### S3.1.1 Pupillometry data acquisition and preprocessing

Pupillometry data were collected using a TRACKPixx3 MRI/MEG (VPixx Technologies, Saint-Bruno, QC Canada), an MRI compatible binocular eye tracker, with sampling rate of 2kHz. Data were preprocessed using the ET-remove artifacts toolbox (https://github.com/EmotionCognitionLab/ET-remove-artifacts), time-shifted by three TRs to align with the hemodynamic response function delay (Murphy et al., 2014), and downsampled to match the temporal resolution of the fMRI data by averaging within each TR (Mather et al., 2020). The measure of interest is percent signal change so that we may measure pupil dilations relative to baseline; therefore, each subject’s pupil dilation time series during PostAr blocks was divided by that subject’s mean pupil dilation during RS0. Three subjects’ data were unfortunately unusable due to technical difficulties during collection procedures, resulting in n = 27 for most pupillometry-related calculations. These three subjects were different from the ones who decided not to participate in the MT-prepared gradient echo scans.

##### S3.1.2 Pupillometry analyses

Pupillometry analyses paralleled some of the state space trajectory analyses. Specifically, we computed the mean pupil dilation changes for state-specific pairwise transitions as a function of condition (active vs sham) to further explore how LC activity may be related to brain state behaviors. Pupil dilation changes as a function of state switching were computed by identifying a switch in subjects’ state sequences, then calculating the difference between the normalized pupil size two TRs before the switch and the first TR after the switch. This calculation was contingent on the subject remaining in the same state for two TRs before or after the identified switch to ensure that they settled into a stable state. We cannot be less stringent with the criterion of remaining in the same state for two TRs before and after the switch because these criteria ensure that changes in pupil size were accompanying specific transitions, and that dilations/contractions from previous or succeeding transitions did not bleed into that calculation. After calculating these differences, the mean of these differences within a subject was found.

Once the mean subject-specific changes in pupil size during state specific transitions were found for both conditions (active and sham), the differences in these values were computed to reveal how pupil dilation changes at specific state transition points might vary due to active squeezing versus sham control. Finally, these differences were Spearman correlated with LC MTC to determine whether LC MTC impacted pupil dilation during switches between HMM-derived latent brain states. Because some subjects’ data were missing due to improper data collection or the aforementioned exclusion criteria, only subjects whose data were accounted for in both the pupillometry and MT-prepared gradient echo datasets were included in these correlations (at most n = 25).

#### S3.2 Pupillometry results and discussion

Above, we saw that the duration spent in S4 once it is entered was correlated with LC MTC, as were changes in transition probabilities between the active and sham condition specifically regarding transitioning from S5→S4. We interpreted these results to mean that LC structure (cell density) plays a meaningful role in the effectiveness of the squeeze manipulation and its consequent effects on brain dynamics, even in the cohort of healthy young adults used here. This is consistent with previous results showing similar relationships between LC structure integrity and cognition in older adults (Hussain et al., 2019; Langley et al., 2020, 2021; Mather & Harley, 2016).

However, because of LC’s small size (∼2mm in diameter), it is difficult to establish whether LC activity itself (measured via the BOLD response) could also drive these state transition characteristics, a problem that is exacerbated by the poor temporal resolution of fMRI. Therefore, as a confirmatory analysis, we turned next to pupillometry data as a proxy for LC activation. As introduced above, pupillary diameter has been established as a viable proxy for LC activity levels (Joshi et al., 2016; Murphy et al., 2014), which makes this measure valuable as a mitigation strategy for the noisiness of LC BOLD signal in fMRI due to the small size of LC.

Importantly, we highlight here that the goal of this analysis was to establish consistency with the results presented above. Unfortunately, due to technical challenges and our stringent criteria for establishing pupil dilation changes as a function of state transition, pupil diameter data was unavailable for some subjects for some analyses (described in detail below). Therefore, we present these results as qualitative confirmation of the patterns identified above, and future studies should seek to remedy the technical challenges we experienced so that a fuller understanding may be gained.

First, we calculated the mean changes in pupil dilation (see **Supplementary Material Section S3.1**) for each pairwise transition between two states, separately as a function of active (**Figure S2A**) and sham (**Figure S2B**) conditions. We then found the difference between them (**Figure S2C**) to highlight distinct pupil size changes resulting from LC activity up-regulation. Finally, the difference in pupil dilation for state-specific transitions across conditions (prior to taking the global average across subjects) was correlated with LC MTC (**Figure S2D**). The criterion that subjects must remain in the same state for two TRs before and after the switch was enforced resulting in different numbers of subjects meeting this standard for a transition and consequent correlation with LC MTC (**Figure S2E**; see also **Methods** in the main text).

**Figure S2.**
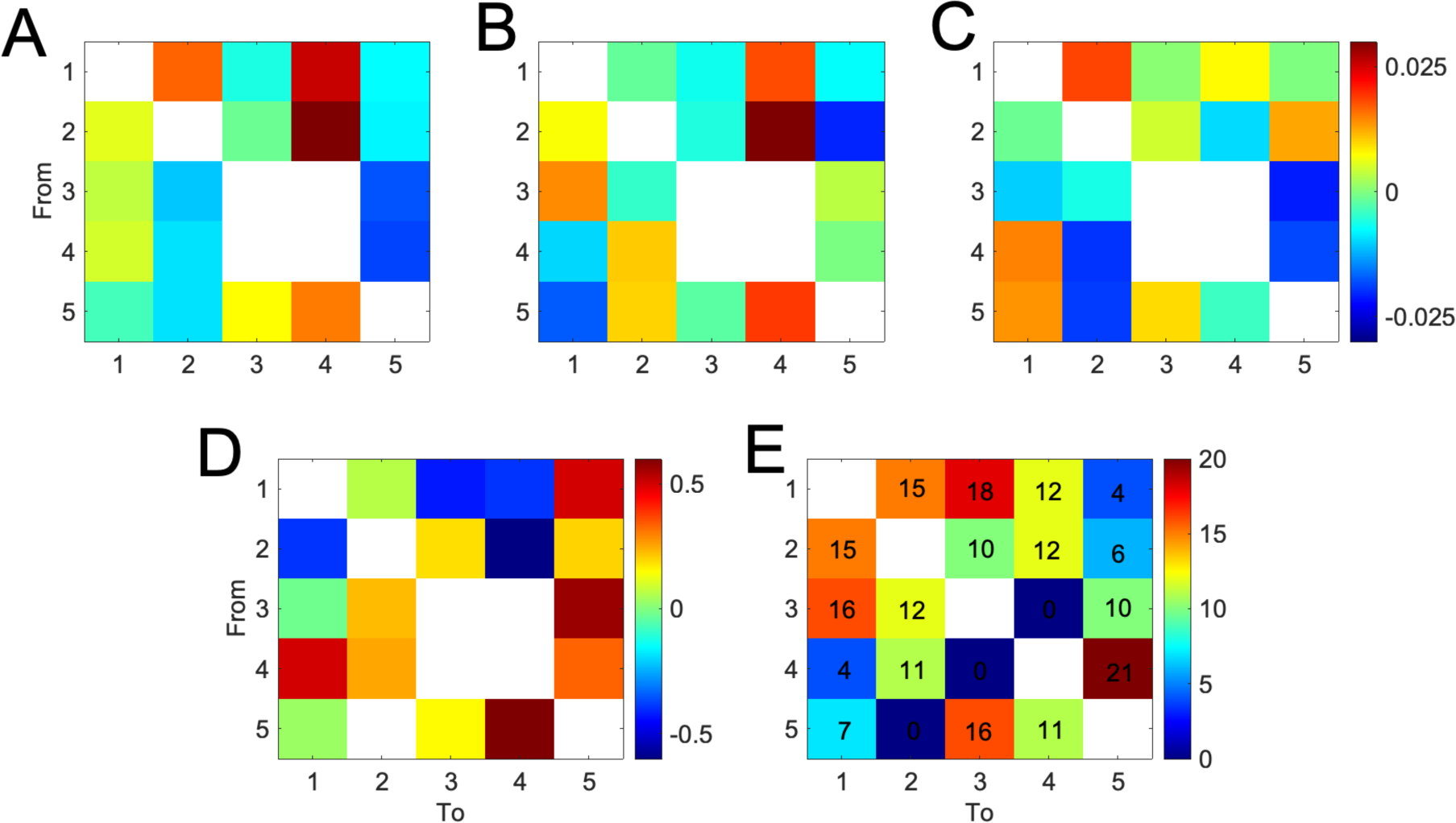
Pupil dilations changes for **(A)** active and **(B)** sham conditions as well as **(C)** the difference between them when subjects transitioned between specific states. **(D)** The difference in pupil dilation changes relative to baseline across conditions was correlated with LC MTC. We computed pupil dilation change for a transition only when the subject remained in the same state for two TRs before and after an identified switch. Not all subjects underwent a switch that met this criterion, so **(E)** shows the number of subjects that experienced each transition. The largest changes in transition-specific pupil dilation to correlate with LC MTC **(D)** occurred for pairwise transitions S1→S5, S3→S5, S4→S1, and S5→S4. Interestingly, three of these four (S1–S5, S4→S1, and S5→S4) coincide precisely with the only three transitions that showed significant changes in relative transition probability matrices (RTPMs; **Figure 8**), and the largest of these (S5→S4) is precisely for the transition that previously showed significant correlation with LC MTC. That is, LC MTC predicted how pupil would react in these specific transitions, and these were also the specific transitions that showed the largest differences between active and sham conditions. White squares occur along the self-transition diagonal as well as marking transitions that never occurred or in **(D)** where no subjects were able to be used for the correlation with LC MTC (see corresponding squares in **(E)**, marked with 0s); see main text for more details.

With the heavy caveat that every ‘square’ in these matrices contains a different number of subjects (**Figure S2E**), we can at least conduct some qualitative explorations. The aspects to focus on are **Figure S2C** and **S2D**, which show the differences between active and sham in transition-specific pupil dilation changes as well as the correlations between these changes and LC MTC across subjects. The differences themselves (**Figure S2C**) may not appear particularly meaningful, but four correlations with LC MTC (**Figure S2D**) stand out: S4→S1, S5→S4, S1→S5, and S3→S5. Notably, three of these four (S4→S1, S5→S4, S1→S5) are the same three that showed significant differences between active and sham conditions in the RTPM analysis, and the largest one (S5→S4) is the specific pairwise transition that showed correlations with LC MTC. This is also consistent with our first observation that S4 average state duration (baseline corrected) significantly correlated with LC MTC as well.

Unfortunately, as mentioned above, the criteria used to define pupil dilation changes did require us to discard a large number of subjects in many of these pairwise state transitions. We hope that this might be remedied by employing a one-hour long paradigm rather than one lasting less than 20 minutes, and by collecting data from more subjects. The chances of switching between states would increase and more subjects would survive the rejection standard consequently increasing the effect size. Nevertheless, despite this challenge we can see qualitative confirmation of the above findings, i.e. that LC MTC is related to the effectiveness of the active squeeze task driving the ease of transitioning into S4 (SN-dominant state) as well as how long subjects persist in occupying S4 once they’ve arrived there (regardless of condition).

Thus, we were only able to show qualitative confirmation of the HMM brain state transition result with our pupillometry data. A likely cause of this result is the stringent criteria we adopted for including a particular instance of pupil dilation changes as a function of a specific pairwise transition. As a result, many subjects were unfortunately excluded as they did not meet this preset criterion of remaining in the same state two TRs before or after a switch. Increasing the number of subjects would boost statistical power to detect this potentially small effect; longer scan durations may also help avoid loss of data usability. Unfortunately, it was not possible to increase the number of subjects or length of scans here, so these possibilities must be left to future studies.

However, there is another possible interpretation of this somewhat weak pupillometry finding. We opted to collect pupillometry data because it has been repeatedly shown that continuous measures of pupil diameter throughout both resting state and task stimuli may index tonic variations in LC BOLD activity, and are less liable to trial-by-trial noise than pupil dilation locked into task-related events (Murphy et al., 2014). Further, electrophysiological studies in monkeys have often shown a reliable relationship between LC activity and changes in pupil diameter due either to spontaneous fluctuations, or to external stimuli (Joshi et al., 2016). However, evidence is now mounting that fluctuations in LC do not necessarily covary with pupil diameter in statistically meaningful ways (Megemont et al., 2022; H. Yang et al., 2021). It therefore seems reasonable to conclude that a lack of strong statistical support from the pupillometry analyses should not color interpretation of our primary network analysis and correlation with LC MTC.

